# Action Potential Thresholds and Excitability from the Geometry of Membrane Potential

**DOI:** 10.64898/2026.08.21.746364

**Authors:** Marco Arieli Herrera-Valdez

## Abstract

A novel mathematical framework to define the threshold of action potentials in excitable cells is presented. Unlike previously applied methods that rely on approximations or bifurcations, the approach focuses on the geometry action potentials. First the changes in concavity directly obtained from a time series of voltages are used to identify action potentials thresholds. This concavity-based framework is subsequently extended to autonomous dynamical systems, enabling the analytical determination of curvature transitions directly from the manifold of inflection points in phase space. The inflection point manifold delineates the boundary of the excitability region, establishing a necessary and sufficient geometric condition where a phase-space trajectory contains an action potential if and only if it intersects this manifold. This geometric framework not only formalises cellular excitability within a single dynamical system but also introduces a measure to compare different electrophysiological phenotypes and stimulus conditions. The measure provides a way to compare the excitabilities of systems that model neurons with different electrophysiological phenotypes and consider different stimulus conditions. The traditionally vague physiological concept of electrical excitability is transformed into a rigorous analytical description by considering the time-dependent curvature of the membrane potential. The criterion is robust across smooth, single compartment models of electrical excitability and can be can be extended to single compartment models in higher dimensions, and multicompartment models as well.

## 1 Introduction

Electrically excitable cells are capable of producing all or none, pulse-shaped deflections in their transmembrane potential, *v* (in mV) (Aidley, 1998). Such pulses named *membrane action potentials (APs)* by Hodgkin and Huxley (1952), participate in different physiological functions that include rapid transmission of signals (Harper and Lawson, 1985), secretion of neurotransmitters and hormones (Dunant, 1994), and regulation of protein expression that supports cellular architecture and plasticity (Desai et al., 1999; Marrone and Petit, 2002). Examples of electrically excitable animal cells include neurons (Cole and Curtis, 1938; Hodgkin and Huxley, 1952), neuroglia (Verkhratsky et al., 2020), striated muscle fibers (Clausen et al., 1998; Stephenson, 1998), cardiomyocytes (Huang and Lei, 2023), pancreatic acinar cells (Petersen, 1982) and *β*-cells (Chay, 1987), and adrenal chromaffin cells (Biales et al., 1976). Of note, there are also electrically excitable cells in plants (Pickard, 1973; Sukhov and Vodeneev, 2009; Wayne, 1993).

A system can be generally defined as excitable if its non-trivial trajectories^1^ display “all-or-none” behaviors (Fig. 1, green and red traces, respectively). Consequently, establishing excitability depends on verifying whether the trajectories in the system can be separated into two groups, APs (the “all”), and non-APs (the “none”). Note that for some cases in neuronal repetitive spiking (Jodkowski et al., 1988) and cardiac oscillations (Campana, 2015; DiFrancesco, 1993; Rasmusson et al., 1990b), if the only attractor is a limit cycle, there could be no partition of the set of trajectories.

**Figure 1.**
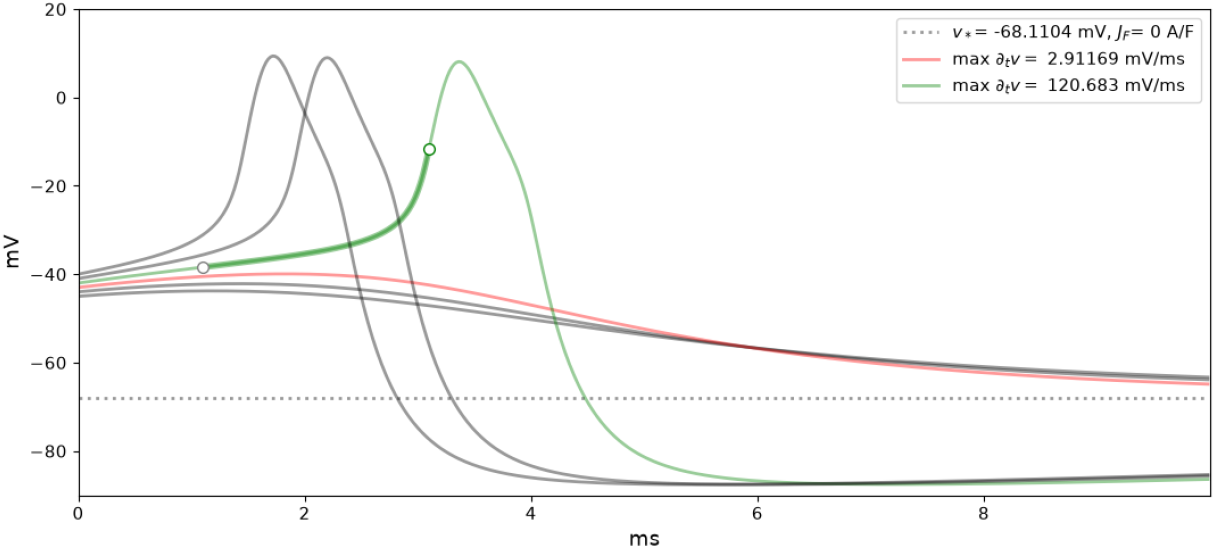
Membrane potential traces from different initial conditions in the absence of current stimulation (*J*_*F*_ = 0). Three APs and three non-APs are shown. The green and red lines show the nearest traces to the transition between APs and non-APs at a resolution of 1 mV, with their maximae ∂_*t*_*v* indicated in the text by color. The increasing and concave-up segment of the AP in green is marked with a thick green line segment, flanked by white circles at the inflection points. The resting potential *v*_∗_ is shown for reference. The traces were obtained from equations (4)-(5) with parameters from Table 1, assuming *w*(0) = *w*_∗_ with *v*(0) ∈ {−47, −39} in steps of 1 mV.

APs and the resulting network activity have been studied as threshold phenomena (Fitz-Hugh, 1955; Gast et al., 2023; Kamaleddin, 2025; Karreman, 1951; Trinh et al., 2023), which places the concept of threshold at the center of analysis. For clarification, a threshold is referred to here as a point along a trajectory (or orbit) at which the behavior of the trajectory is fundamentally altered.

**Table 1.** Parameters for the model. All the rate amplitudes for the currents are maximal membrane conductances normalized by the product of *v*_T_ and the membrane capacitance (Herrera-Valdez, 2018).

| Parameter | Value/Range | Units | Meaning |
| --- | --- | --- | --- |
| $C_m$ | 20 | pF | Membrane capacitance |
| $J_F$ | $\geq 0$ | A/F | Input stimulus current (pA) normalized by membrane capacitance (pF) |
| **Voltages** |  |  |  |
| $v_T$ | 26.7 | mV | Thermal potential. |
| $v_K$ | -70 | mV | Nernst potential for $K^+$ |
| $v_{Na}$ | 60 | mV | Nernst potential for $Na^+$ |
| $v_{NK}$ | -60 | mV | Na-K ATPase reversal potential |
| $v_w$ | 0 | mV | Half-activation potential for KD channels |
| $v_m$ | -20 | mV | Half-activation potential for NaT channels |
| **Rates** |  |  |  |
| $a_{NK}$ | 0.25 | 1/ms | Amplitude for the Na-K ATPase current |
| $a_{KD}$ | 13, 26 | 1/ms | Amplitude for the current mediated by KD channels |
| $a_{NT}$ | 2.5 | 1/ms | Amplitude for the current mediated by NaT channels |
| $a_w$ | 0.1 | 1/ms | Activation rate for KD channels |
| **Gains and biases** |  |  |  |
| $\kappa$ | 0 | – | Exponent that allows the possibility of observing logistic behavior in $w$ for fixed $v$ |
| $b_w$ | 0.65 | – | Activation bias for KD channels |
| $s_w$ | $3/v_T$ | 1/mV | Gain for KD channel activation, normalized by $v_T$ |
| $s_m$ | $5/v_T$ | 1/mV | Gain for NaT channel activation |

The voltage at which an AP begins can be thought of as a point of “kinetic commitment” after which *v*(*t*) increases and accelerates. The initiation of an action potential thus represents a threshold marking the onset of the upstroke phase, during which *v*(*t*) rises toward its peak (∂_*t*_*v* = 0), first accelerating and subsequently decelerating. Geometrically, this means that the graph of *v*(*t*) increases, becomes concave-up, and then changes concavity before the peak (Fig. 1, green trace). In contrast, trajectories that do not describe APs (Fig. 1, red trace and gray traces below) do not exhibit an a up-down change in curvature with respect to time before they reach their peak.

The accelerating portion of the AP upstroke is thus a defining feature of the AP and it is bounded by two inflection points that can be regarded as thresholds (Spivak, 2018). The first inflection point is where *v*(*t*) “lifts-off” and corresponds to a minimum for the rate of increase of *v*(*t*). The second inflection point occurs at the maximum rate of increase for *v*. Both of these points bound the accelerating portion of the AP upstroke and *v*(*t*) changes concavity there. Then, ∂_*t*_*v >* 0 and 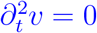 at *v* = *v*_LO_ and *v* = *v*_MR_, and *v*(*t*) is concave-up (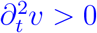) between *v*_LO_ and *v*_MR_ (Fig. 1 white dots and thick green line). The times *t*_LO_ and *t*_MR_ at which the “lift-off” and maximum rate of increase occur can then be regarded as threshold times, and *v*_LO_ = *v*(*t*_LO_) and *v*_MR_ = *v*(*t*_MR_) would then be threshold voltages.

The concave-up segment of the upstroke 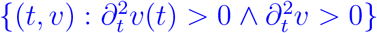 (Fig. 1, green, thick segment) is thus a defining feature that seems to be necessary and sufficient to identify an AP.

To address these geometric and physiological challenges, this paper seeks to:

- Identify whether action potentials can be detected by finding *v*_*LO*_ and *v*_*MR*_ in voltage time series (Section 2).
- Calculate analytically the concavity and inflection points for *v*(*t*) in autonomous dynamical systems (ADS, Section 3.2.1) to find the inflection points and formalise the concept of AP thresholds (3.2.2).
- Formulate a phase-plane criterion for the occurrence of APs to formalise the definition of excitability as an “all-or-none” (APs and non-APs) partition of phase-space trajectories (Section 3.3).
- Examine how these threshold manifolds shift under different biophysical modulations (Section 3.4).
- Define a quantitative measure of excitability (Section 3.5) to compare different phenotypes and stimulus conditions, and describe the excitability of model membranes across various electrophysiological profiles (Section 3.5.1).
- Generalize these curvature and threshold calculations to higher-dimensional and multi-compartment models (Section 3.6 and Section 3.7).

## 2 Concavity in randomly fluctuating voltage traces: a quick test by application

Assume *v* is recorded at times *t*_0_, *t*_2_, … *t*_*n*_. The regions where the first and second derivative of *v* are positive can be found by approximating the values ∂_*t*_*v* and 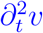 by secant lines obtained directly from *v*(*t*_0_), …, *v*(*t*_*n*_). Explicitly, a time series *D*(*t*_0_), …, *D*(*t*_*n*_) approximating ∂_*t*_*v* at *t*_0_, …, *t*_*n*_ can be calculated by letting

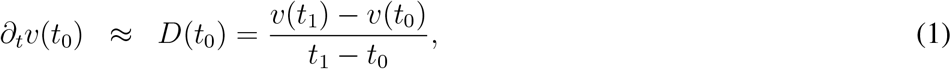

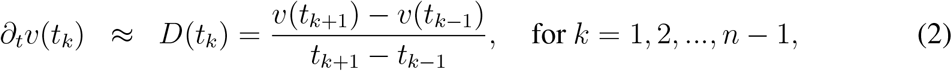

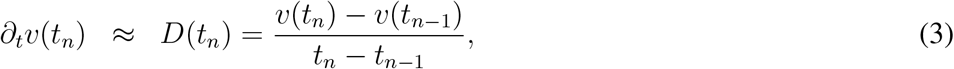

A similar calculation can be performed to obtain a time series *D*^2^(*t*_0_), …, *D*^2^(*t*_*n*_) approximating 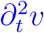 at *t*_0_, …, *t*_*n*_.

To find the AP upstroke inflection points (AP thresholds, Fig. 2, yellow and orange dots), (1) Extract the collection *C*_1_ of indices *i* where *v* is increasing and concave-up (condition 1: *D*(*t*_*i*_) *>* 0 and *D*^2^(*t*_*i*_) *>* 0). (2) To eliminate false-positive segments where condition 1 is fulfilled and in consideration that APs should last at least half a ms, identify pairs of consecutive indices from *C*_1_ that correspond to time segments larger than half a millisecond (*e*.*g*.a minimum difference of 20 points at a sampling rate of 40 kHz). Each such pair contains the indices that correspond to a maximum rate inflection point, and the next minimum rate inflection point, respectively. Put all the indices for the maximum rates in one array, say *I*_*M*_ and all the indices for the minimum rates in another array, say *I*_*m*_. (3) The index of the first inflection point of the first AP is the first of the consecutive indices in *C*_1_ that precedes the first index in *I*_*M*_ . Add this index to *I*_*m*_. (4) The index of the last inflection point of the last AP is the last of the consecutive indices in *C*_1_ that are larger than the last index in *I*_*m*_. Add this index to *I*_*M*_ .

**Figure 2.**
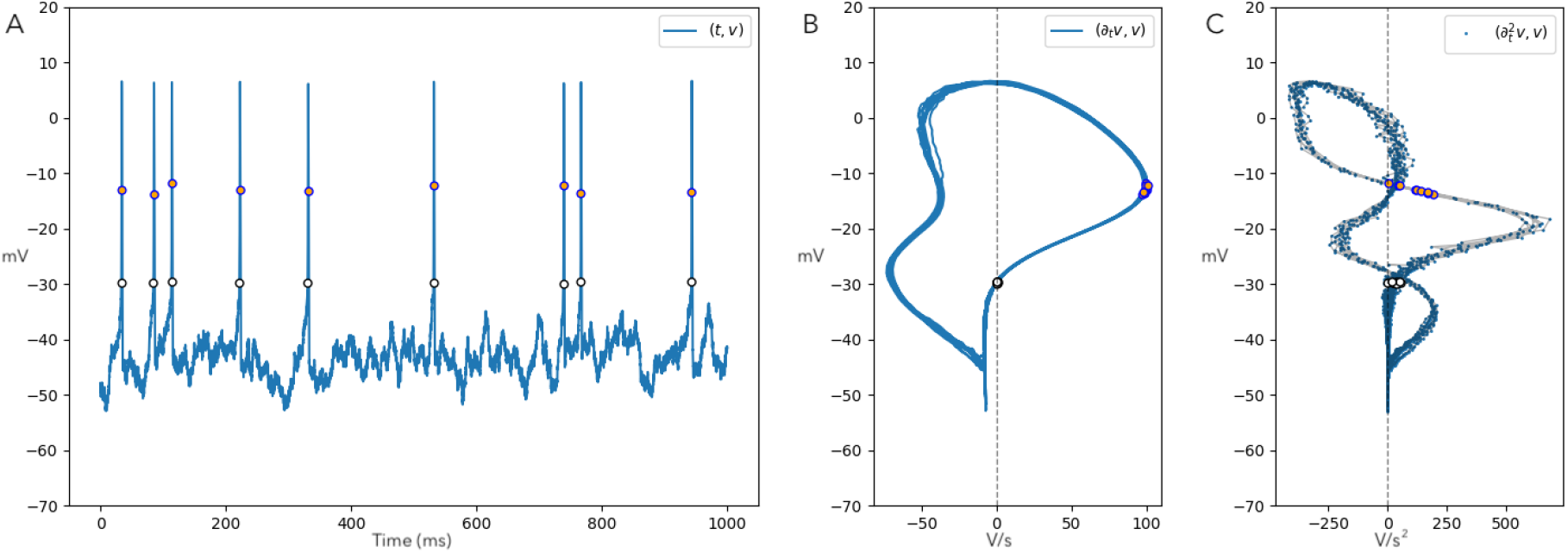
Detection of action potential times from voltage time series. The upstroke inflection points (yellow and orange circles in each plot, respectively) are found using equations (1)-(3). (A) *v* as a function of time; (B) y (C) phase planes (∂_*t*_*v, v*) and 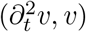, respectively. The time traces were generated from numerical solutions of model in equations (4)-(5) forced with a Gaussian process.

From here on, a threshold is referred to as a point that is crossed by a trajectory, such that the trajectory displays a qualitatively important property not displayed before crossing the threshold.

## 3 AP thresholds and excitability in biophysical, 2-dimensional models of membrane potential

As seen in the last section, finding the inflection points of *v* with respect to time allows detecting APs with a precise, mathematical criterion involving the first and second time-derivatives of *v*. Now let us explore the geometry of AP trajectories in models based on ADS, which reproduce dynamics observed in “box-shaped” current clamp experiments. One important construction assumption in these models is that the biophysical properties of a neuron (*e*.*g*. channel expression) do not change throughout the duration of a current-clamp experiment. As a consequence, these models are families of ADS in which the functional form of the equations is the same for all systems, and the external input parameter is allowed to change while all other parameters remain constant (one autonomous system for each value of the stimulus)^2^. Can we use the concavity criterion from last section to find AP-thresholds and regions in phase space where trajectories are APs? Is it possible to calculate the curvature and inflection points analytically in these models? If that was the case, it should also be possible to provide, by extension, a precise definition for an excitable ADS based on identifying trajectories containing APs and trajectories without them.

### 3.1 Membrane dynamics

The biophysical models of neuronal membrane potential *v* of the lowest dimension that can be constructed with continuous, autonomous equations are 2-dimensional (Av-Ron et al., 1991; Fohlmeister et al., 1990). Such models can be constructed by assuming that the change in *v* with respect to time is a consequence of transmembrane K^+^and Na^+^fluxes mediated by Na^+^-K^+^ ATPases (*J*_NK_), voltage-gated delayed rectifier K^+^-channels (*J*_KD_), and voltage-gated inactivating Na^+^ channels (*J*_NK_) (Herrera-Valdez, 2018), all normalized by the membrane capacitance *C*_*m*_ (Herrera-Valdez, 2020), in units of A/F. The second variable *w* represents the proportion of activated voltage-gated K-channels (Herrera-Valdez and Lega, 2011; Rinzel, 1985). The dynamics can be explicitly described by equations of the form

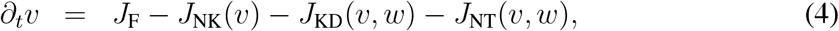

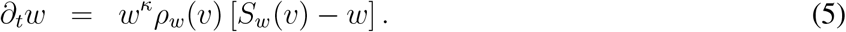

with *w* ∈ [0, 1] and *v* ∈ [*v*_K_, *v*_Na_], with *v*_K_ and *v*_Na_ representing the Nernst potentials for K^+^ and Na^+^, respectively (Herrera-Valdez, 2020). All the systems taken into consideration from here on will be defined within the domain *D* = [0, 1] *×* [*v*_K_, *v*_Na_]. *J*_F_ (also in A/F) represents the amplitude of the current stimulus after normalization by *C*_*m*_. Note that *w* also symbolizes the proportion of inactivated Na^+^-channels, so 1 − *w* represents the proportion of non-inactivated Na^+^channels. The term *w*^*κ*^ yields solutions for *w* with logistic shape if *κ >* 0 and *v* is fixed, as shown in experimentally recorded currents in voltage-clamps (see Fig. 3 in the classical paper by Hodgkin and Huxley (1952) and in Tsunoda and Salkoff (1995)).

**Figure 3.**
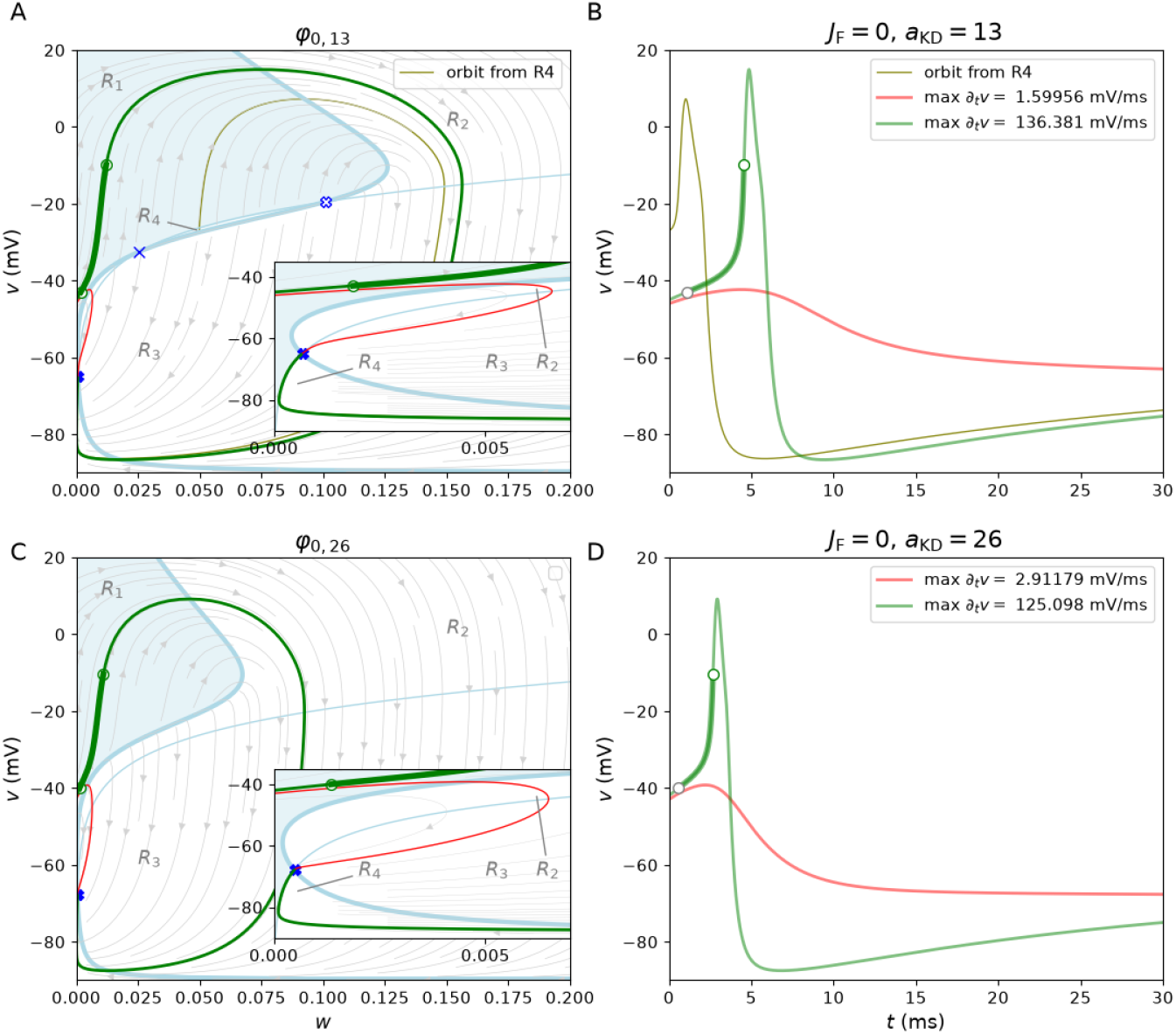
Two phase planes with their corresponding (*t, v*) plots illustrating the two topologically different configurations of dynamical systems from equations (4)-(13) for *J*_*F*_ = 0. The green and red lines show an AP a non-AP, respectively. The inflection points where *v* = *v*_LO_ and *v* = *v*_MR_ in the AP (calculated as in Section 2) are shown in white. The phase plane thin and thick blue lines show the *w*- and *v*-nullclines, respectively. The stream lines formed by the vector field are shown in light gray. The phase plane region *R*_1_ is shaded in light blue and *R*_2_, *R*_3_, and *R*_4_ are annotated for reference. Insets show the dynamics near the attractor fixed point. (A-B) System with 3 fixed points (*φ*_13,0_): an attractor node, a saddle, and repulsive node, respectively marked with a thick blue cross, a thin blue cross, and a thick white cross. The olive line shows an AP that starts in region *R*_4_, near the separatrix segment formed by the portion of the *v*-nullcline between the saddle and the repulsive node. (C-D) System with 1 an attractor node fixed point (*φ*_26,0_).

Explicitly, the currents can be written in general terms as

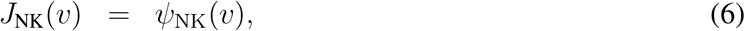

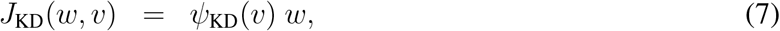

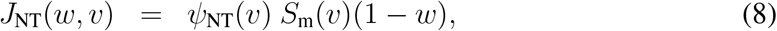

with

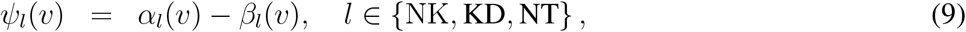

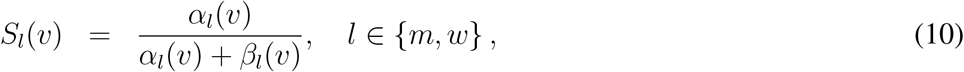

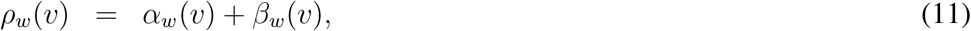

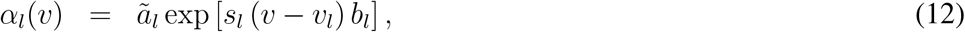

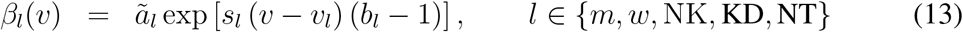

The terms *α*_*l*_, *β*_*l*_, and *ψ*_*l*_ for *l* ∈ {NK, KD, NT} respectively represent the forward and backward flows and the driving forces for the transmembrane currents^3^ (Herrera-Valdez, 2018). *α*_*l*_ and *β*_*l*_, *l* ∈ {*w, m*} represent forward and backward rates of activation for the voltage-gated channels. The amplitude term ã_*l*_ is in units of 1/ms if *l* ∈ {*m, w*}. If *l* ∈ {NK, KD, NT}, ã_*l*_ = *N*_*l*_*A*_*l*_*/C*_*m*_ is in units of A/F with *A*_*l*_ in pA, *C*_*m*_ in pF, *N*_*l*_ representing the number of transmembrane proteins mediating the mechanism *l, v*_*l*_ is the reversal potential, *b*_*l*_ the bias (rectification) of the transmembrane motion of ions in the energetically favorable direction for *l* ∈ {NK, KD, NT}.

Sacrifying the first principles approach, rectification, and the nonlinearities in the thermodynamical model for the sake of numerical speed and notational simplicity, conductance-based models (Hodgkin and Huxley, 1952) can be obtained from equation (9) by calculating a linear approximation around the reversal potential *v*_*l*_ (Herrera-Valdez, 2018) that yields

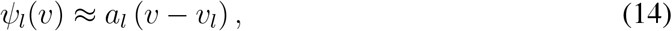

with 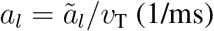.

From a dynamical systems point of view, *v* can be regarded as a fast and self-amplifying variable that positively feeds into *w*. In turn, *w* has a slower evolution and negatively feeds back into *v*. The slower dynamics of *w* allow some the trajectories to increase abruptly in *v* before *w* makes it decrease. That is the set of orbits where APs can be found.

#### Model characteristics and constrains for parameters

The APs should be such that 100 ≤ max {∂_*t*_*v*} ≤ 300 V/s, last between 1 and 3 milliseconds, with amplitudes between 80 and 100 mV, with a rheobase^4^ of approximately 20 pA (*J*_*F*_ ≈ 1 A/F if the membrane capacitance is 20 pF) for the most excitable electrophysiological phenotypes.

### 3.2 Geometry under lying thresholding and excitability from an analytical perspective

To start studying the geometry of the system, rewrite equations (4)-(14) as

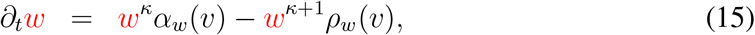

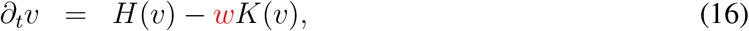

with

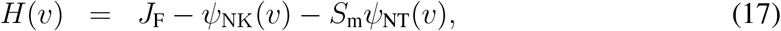

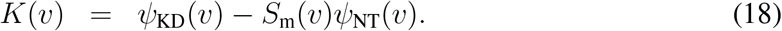

From there, it is possible to analytically obtain expressions for the nullclines, curvature, and determine the stability and type of the fixed points (see Appendix, Section A and Section D).

First, consider the family of dynamical systems φ with evolution given by equations (15)-(18) and parameters as in Table 1. Assume that the dynamics for K-channel activation and the Na-channel inactivation are linear (*κ* = 0) and that there is no stimulus current (*J*_F_ = 0, Fig. 3). The system can then be studied in two possible configurations: φ_13,0_ obtained by setting *a*_KD_ = 13, with three fixed points (Fig. 3A-B), and φ_26,0_, representing a neuron with twice as many K-channels, *a*_KD_ = 26 (Fig. 3C-D), but identical voltage-dependent gating properties for all channels. Of note, the phase planes for both dynamical systems φ_13,0_ and φ_26,0_ are divided by the nullclines into four regions labeled *R*_1_,…,*R*_4_, defined by alternating the signs of ∂_*t*_*v* and ∂_*t*_*w* (Table 2 and Fig. 3).

**Table 2.** Regions of the phase plane defined by the nullclines in equations (40)-(41). The region *R*_1_ contains the upstokes of all APs (Fig. 3).

| Region | Description | $\partial_t w$ | $\partial_t v$ | AP segments contained |
| --- | --- | --- | --- | --- |
| $R_1$ | $\left\{ (w, v) : 0 < w < \min \left( \frac{H(v)}{K(v)}, \frac{\alpha(v)}{\rho(v)} \right) \right\}$ | + | + | AP upstroke, return to resting potential |
| $R_2$ | $\left\{ (w, v) : \frac{H(v)}{K(v)} < w < \frac{\alpha(v)}{\rho(v)} \right\}$ | + | - | Initial Downstroke |
| $R_3$ | $\left\{ (w, v) : \max \left( \frac{H(v)}{K(v)}, \frac{\alpha(v)}{\rho(v)} \right) < w < 1 \right\}$ | - | - | Afterhyperpolarization |
| $R_4$ | $\left\{ (w, v) : \frac{H(v)}{K(v)} > w > \frac{\alpha(v)}{\rho(v)} \right\}$ | - | + | Initial repolarization and upstroke if the system has 3 fixed points (central region between saddle and repulsor fixed point in Fig. 3) |

In both configurations φ_13,0_ and φ_26,0_ of the system, it is possible to find APs (Fig. 3, green lines) starting near a non-AP within an arbitrary distance (*e*.*g*. 1 mV, Fig. 3, red lines). Therefore, the threshold phenomenon mentioned earlier is present in these phase planes. Further, all the upstrokes for APs are found in *R*_1_, the region where both variables increase (∂_*t*_*w >* 0 and ∂_*t*_*v >* 0). Most of the discussion will focus on this region.

#### 3.2.1 Analytical calculation of curvature

The curvature for *v*(*t*) can be obtained from equation (16) by calculating

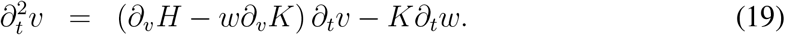

The function 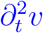 yields a single value of curvature for each state (*w, v*). In particular, the points (*w, v*) for which 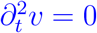 are the inflection points of the whole system (Fig 4, brown lines). Explicitly,

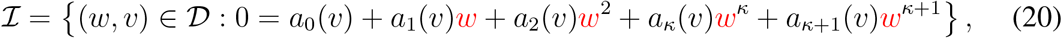

where

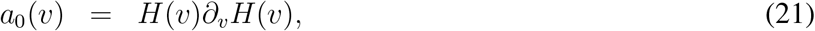

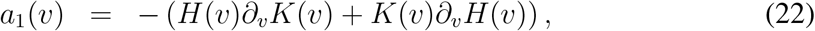

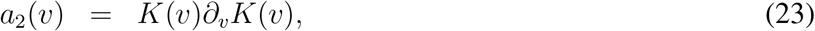

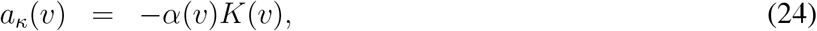

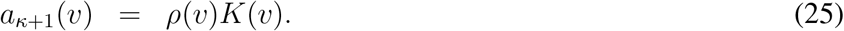

**Figure 4.**
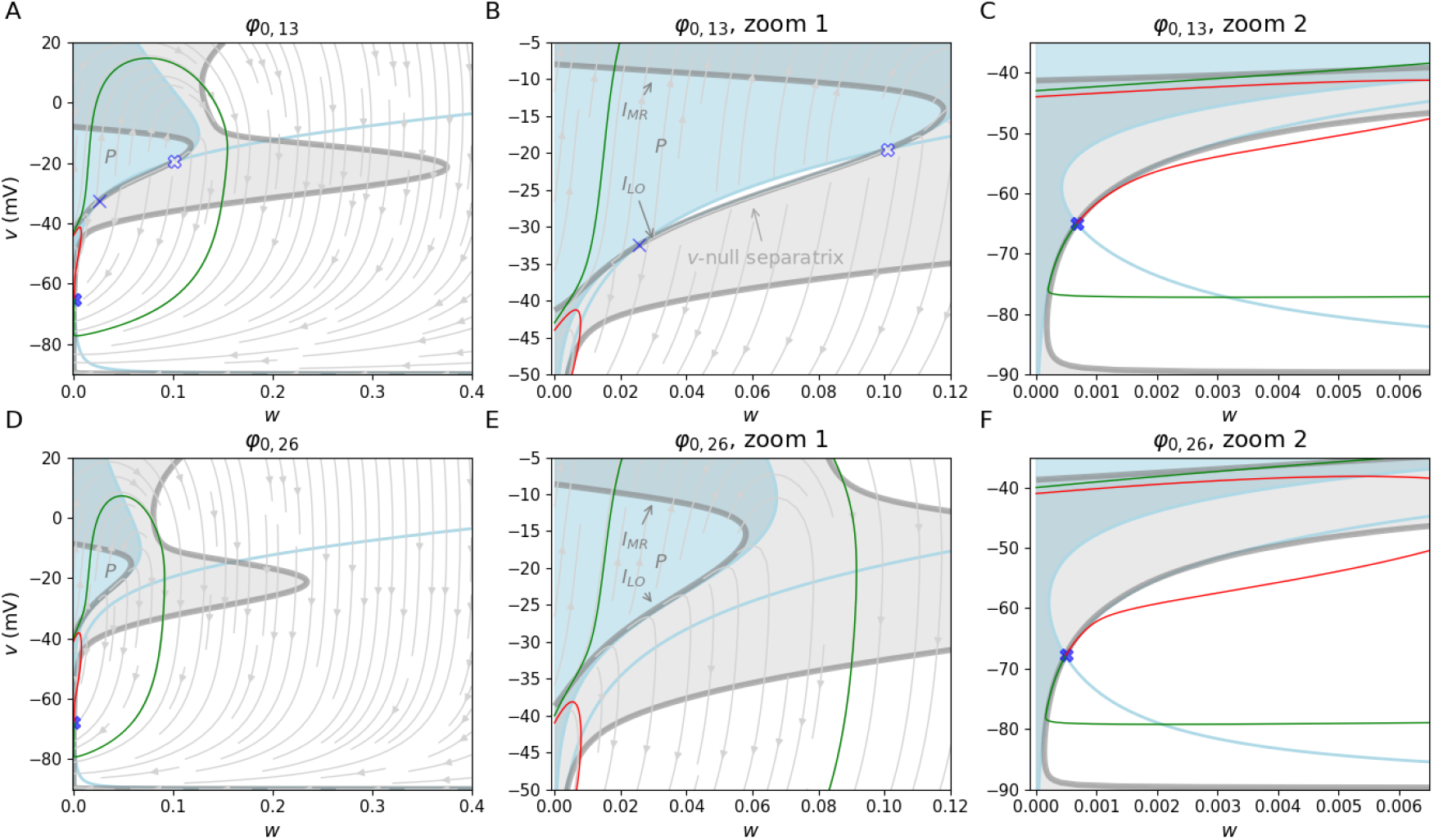
Curvature changes in the phase plane for *J*_*F*_ = 0 (increasing *a*_KD_ while leaving *a*_NT_ fixed. The light blue lines show the *v*- and *w*-nullclines, respectively. The green and red trajectories illustrate APs and non-AP, respectively. The two components of ℐ are shown in thick gray lines. The white and light gray areas are zones where *v*(*t*) is concave-up, and concave-down, respectively. *R*_1_ is shaded in light blue. The ℐ_LO_ and ℐ_MR_ segments of the curve ℐ_*θ*_ ⊂ *R*_1_ are annotated in B and E.(A-C) System with three fixed points (*φ*_13,0_). Note the separatrix segment part of the *v*-nullcline in panel B. (D-F) System with 1 fixed point (*φ*_26,0_). Panels B-C show zoomed versions of A. The same occurs for panels E-F with respect to D. *U*_1_ is annotated in A,B,D,E.

If all parameters are fixed, ℐis a 0-level set in the phase plane, and each point in ℐ corresponds with a change in concavity for *v*(*t*).

##### Special case: quadratic solutions for

*κ* = 0. Note that the *w*-values of the curve ℐ be expressed explicitly as a function of *v* when *κ* = 0. Explicitly, the set of inflection points is given by

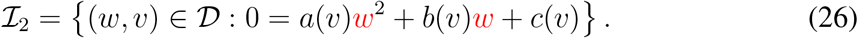

In that case the coefficients transform into

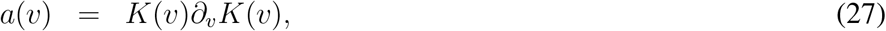

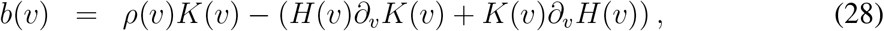

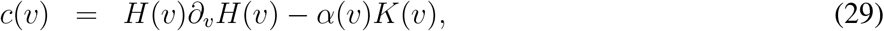

and the solutions in that case would be

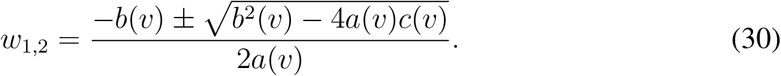

#### 3.2.2 Inflection points for *J*_*F*_ = 0

The curve *I* has two connected components in *D* for both systems φ_13,0_ and φ_0,26,0_ (Fig. 4, brown lines). The right-most component is crossed mostly by segments in the repolarization phase APs, Fig. 4A,D), or by orbit segments traversing through *R*_4_ and *R*_1_ in which *v* increases while going toward the attractor point from below (Fig. 4C,F).

The left-most component of ℐseparates *R*_1_ vertically into three regions: two where *v*(*t*) is concave-down (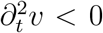, Fig. 4, gray patches), flanking a set where *v*(*t*) is concave-up (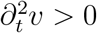, Fig. 4, white patches).

Since *R*_1_ contains all the AP upstroke segments, the set *I*_*θ*_ = *I R*_1_, is the set of inflection points in the AP-upstrokes. ℐ_*θ*_ is a continuous curve formed by the union of a lower segment ℐ_LO_ and an upper segment ℐ_MR_ (Fig. 4B,E). *I*_LO_ is made of points where ∂_*t*_*v >* 0 and reaches a minimum. The points where trajectories cross ℐ_LO_ are the points where *v*(*t*) starts to accelerate as it changes from concave-down to concave-up (Fig. 4B,E, and C,F).

The upper segment ℐ_MR_ contains points at which trajectories reach their maximum rate of depolarization and *v*(*t*) changes from concave-up to concave-down on its way to the AP peak. Importantly, not all the AP trajectories start with *v*(*t*) concave-down (see trajectories starting above ℐ_LO_ near the *v*-axis). Nevertheless, all AP trajectories cross ℐ_MR_. On the other hand, non-AP trajectories do not cross ℐ_*θ*_.

### 3.3 Excitability in a single dynamical system

As already mentioned, a definitive characteristic of an AP is a simultaneous increase in *v*(*t*) and its time derivative (∂_*t*_*v >* 0 and 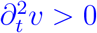). The increase in *v* causes the transmembrane currents to be such that

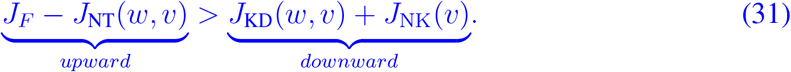

The “acceleration” causes the difference between the left and right sides of inequality (31) to increase (“upward” push minus “downward” push, for *J*_*F*_ ≥ 0 and *v* ≥ *v*_NK_). An AP trajectory that crosses ℐ_*θ*_ twice starts accelerating at (*w*_LO_, *v*_LO_), the point where the trajectory crosses ℐ_LO_, and continues until the maximum value of ∂_*t*_*v* is reached at (*w*_MR_, *v*_MR_), the point where the trajectory crosses ℐ_MR_. After that, the difference between the “up-pushing” and “down-pushing” currents starts to decrease but remains positive.

#### There is a region only crossed by APs that contains all the AP upstrokes

Consider the set

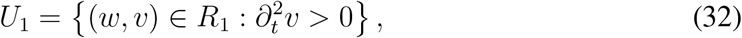

(Fig. 4 A-B, C-D). Note that the set of upstroke inflection points ℐ_*θ*_ is the boundary of *U*_1_. Also notice that all the orbits with upstokes pass through *U*_1_ before reaching their maximum ∂_*t*_*v* on their way to their peak. That is, all the trajectories passing through *U*_1_ contain at least one AP, and the trajectories that do not cross *U*_1_ do not contain APs. Using this criterion, trajectories containing an upstroke with a concave-up segment in *v*(*t*) (passing through a region *U*_1_) will be referred to as AP orbits or AP trajectories. This means that AP orbits cross ℐ_*θ*_ at least once at ℐ_MR_, and possibly at ℐ_LO_.

#### A definition of excitability in single dynamical systems

A dynamical system can be called “excitable” if it contains AP orbits, and non-AP orbits that do not start in a fixed point. A dynamical system φ has AP orbits if and only if it has a region *U*_1_. As a consequence, *U*_1_ will be called from here on the *excitability set*, and a dynamical system will be regarded as excitable if it has a region like *U*_1_. The boundary of inflection points within *R*_1_ can then be regarded as a threshold set. As mentioned before, an autonomous model of membrane potential is a family of dynamical systems in which the parameter for current stimulus could take different values, but all other parameters and dynamics are identical (one ADS for each value of *J*_F_). By extension, an autonomous model of membrane potential can be called excitable if at least one of its ADS is excitable.

#### Excitable *vs*. Excited

Dynamical systems possessing a *unique* limit cycle attractor (repetitive firing) cannot be called excitable because all its nontrivial trajectories converge to the same limit cycle attractor. This means that the trajectories cannot be partitioned into two sets containing AP and non-trivial non-AP orbits. Such systems will be called *excited*, not excitable. Using a similar reasoning, bistable systems with one limit cycle attractor and one attracting fixed point are excitable.

Note that an AP orbit may contain more than one AP, as would be the case for an AP orbit that converges to a limit cycle (Section C in the Appendix). In that case the orbit contains an infinite number of APs.

It is important to remark that some AP orbits converge to a fixed point before covering the entire voltage range of its initial ascent, as it happens in a depolarization block. The resulting waveform can be regarded as containing an incomplete, or partially repolarized AP.

The requirement that an orbit must exhibit a concave-up segment in *v*(*t*) during its upstroke to qualify as an AP is clearly illustrated by the fact that *not all trajectories initiating in R*_1_ *become action potentials* (Fig. 4, red traces). Trajectories of this kind start below ℐ_LO_, never cross ℐ_*θ*_ or touch the region *U*_1_, and are such that *v*(*t*) is increasing and concave-down until they reach their peak at the *v*-nullcline, and decrease on their way to the attractor (Fig. 4, green traces). For a concrete example, there is a region *U*_1_ in φ_13,0_ and in φ_26,0_, and both systems have an attractor node to which all non-trivial trajectories converge (Appendix, Section C). Also, *not all the nontrivial trajectories pass through U*_1_, in both systems. The set *T*_*P*_ of trajectories that passes through *U*_1_, and its complement *D \ T*_*P*_, divide *D* into two sets of trajectories that are either APs and non-APs (trivial and non-trivial), respectively. This means that the systems φ_13,0_ and φ_26,0_ are both excitable.

### 3.4 AP thresholds and biophysics

From current clamp experiments we know that increasing the stimulus amplitude is likely to increase the neuron’s capability of producing APs (Carter and Bean, 2009; Mainen and Sejnowski, 1995). From modeling based on ADS we know that increasing the stimulus amplitude changes the number or the types of fixed points, and their attractivity (Herrera-Valdez, 2012, 2018). In addition, changing biophysical parameters like the number of potassium channels (Zeberg et al., 2015) also has effects on the geometry of the phase space, and in general, on the bifurcation structure in models.

#### The lift-off threshold *v*_LO_ is raised by K-channel activation and by Na-channel inactivation

Taking the two systems φ_13,0_ and φ_26,0_ for illustration, the *v*-values of ℐ_LO_ (Fig. 4A-B,D-E) increase as *w* increases. That is, ℐ_LO_ can be thought of as an increasing function of *w*, which means that the lift-off voltage for APs increases with the activation of *K*-channels.

Similarly, larger proportions of inactivated Na-channels yield higher lift-off *v*_LO_ thresholds, in agreement with the intuition that the upstroke is more likely to occur if more Na-channels are available to activate.

#### The maximum rate threshold *v*_MR_ decreases if there are more activated K^+^channels

Follow the upper portion ℐ_MR_ of the curve ℐ_*θ*_ from left to right (Fig. 4A-B,D-E). The voltage thresholds *v*_MR_ from points in *I*_MR_ are lower for larger values of *w*, for φ_13,0_ and φ_26,0_. In other words, larger proportions of activated K-channels yield lower maximum depolarization rates.

Similarly, if the proportion of non-inactivated Na^+^channels decreases, then the voltage threshold for the maximum rate *v*_MR_ increases. This is in agreement with the intuition that increasing Na^+^channel inactivation should result in slower AP upstrokes.

Now consider following ℐ_LO_ in φ_13,0_ (3 fixed points), from left to right in the phase plane. Note that non-AP trajectories starting at *R*_1_ reach the *v*-nullcline before reaching the (middle) saddle-node fixed point. That is, the *v*-peaks of non-APs starting at *R*_1_ are bounded by above by the *v*-coordinate of the saddle fixed point. Moving to the right in the phase plane, non-AP trajectories can only be found below the portion of the *v*-nullcline located between the saddle and the repeller node fixed points (Fig. 4A-C) where the *w*-nullcline becomes the boundary for *R*_1_). Therefore, the set of non-APs in φ_13,0_ to the right of the saddle point is bound from above by the attracting portion of the *v*-nullcline saddle-node fixed point, as already reported by other researchers (Izhikevich, 2007), (Fig. 4A-C).

In contrast, in φ_26,0_ (1 fixed point), the non-AP trajectory starting in *R*_1_ with the highest local maximum crosses the *v*-nullcline near the point where ℐ_LO_ meets ℐ_MR_ (Fig. 4D-E).

### 3.5 A measure of excitability

Can we quantitatively assess how excitable is a dynamical system? If so, can that quantification be used to compare excitabilities in pairs of dynamical systems?

One way to do so is to measure the size of the region *U*_1_ within the domain *D*. To do so,

Let

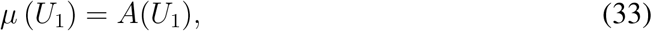

where *A*(·) represents the area (Fig. 5).

**Figure 5.**
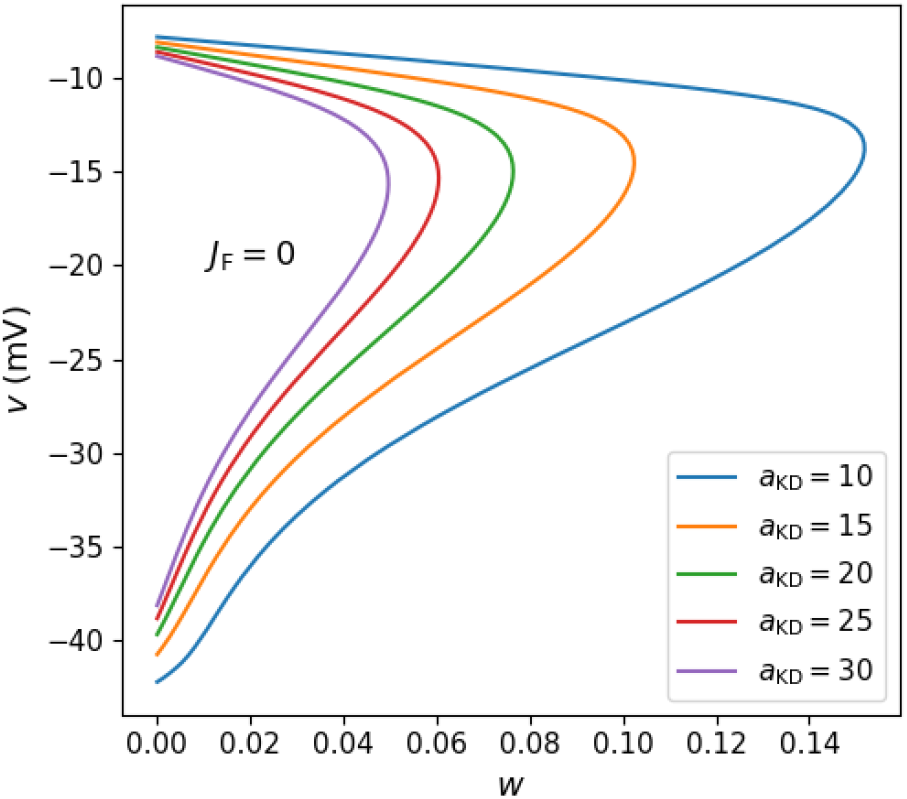
Progression of ℐ_*θ*_ for *a*_KD_ ∈ {10, 15, 20, 25, 30}, and *J*_*F*_ = 0. All other parameters fixed.

For computational purposes, such a measure could be approximated by setting up a grid on *D* with a fine mesh, counting the number of points in *U*_1_ multiplied by the area of one rectangle in the grid. The process can be repeated by increasing the number of points in the mesh for *D*, and stopped when the difference between two values of *µ*(*U*_1_) is less than some given tolerance.

#### 3.5.1 Excitability for electrophysiological phenotypes and input currents

To develop some intuition about the change in the manifold ℐ_*θ*_ for different electrophysiological phenotypes, consider first increasing the number of K^+^channels, while keeping *J*_F_ = 0 and fixing the number of Na^+^channels so that the maximum ∂_*t*_*v* remains within the correct range. Is it reasonable to expect AP thresholds to change? The answer can be obtained by observing the changes in the threshold manifold ℐ_*θ*_. As the number of K^+^channels increases, the curves describing ℐ_*θ*_ keep their shape, but decrease in length and the areas they enclose (*i*.*e. U*_1_) form a nested set of decreasing size (Fig. 5).

Now let us use the measure *µ* to assess the excitability of a model neuron for different electrophysiological profiles and different stimulus amplitudes. Comparing different values of the measure *µ*(*U*_1_) for increasing values of *J*_F_, as long as *J*_F_ is under the rheobase, yields a monotonic, increasing function for each electrophysiological profile. This result is in line with the intuition that larger injected currents yield more excitable cells (Fig. 6).

**Figure 6.**
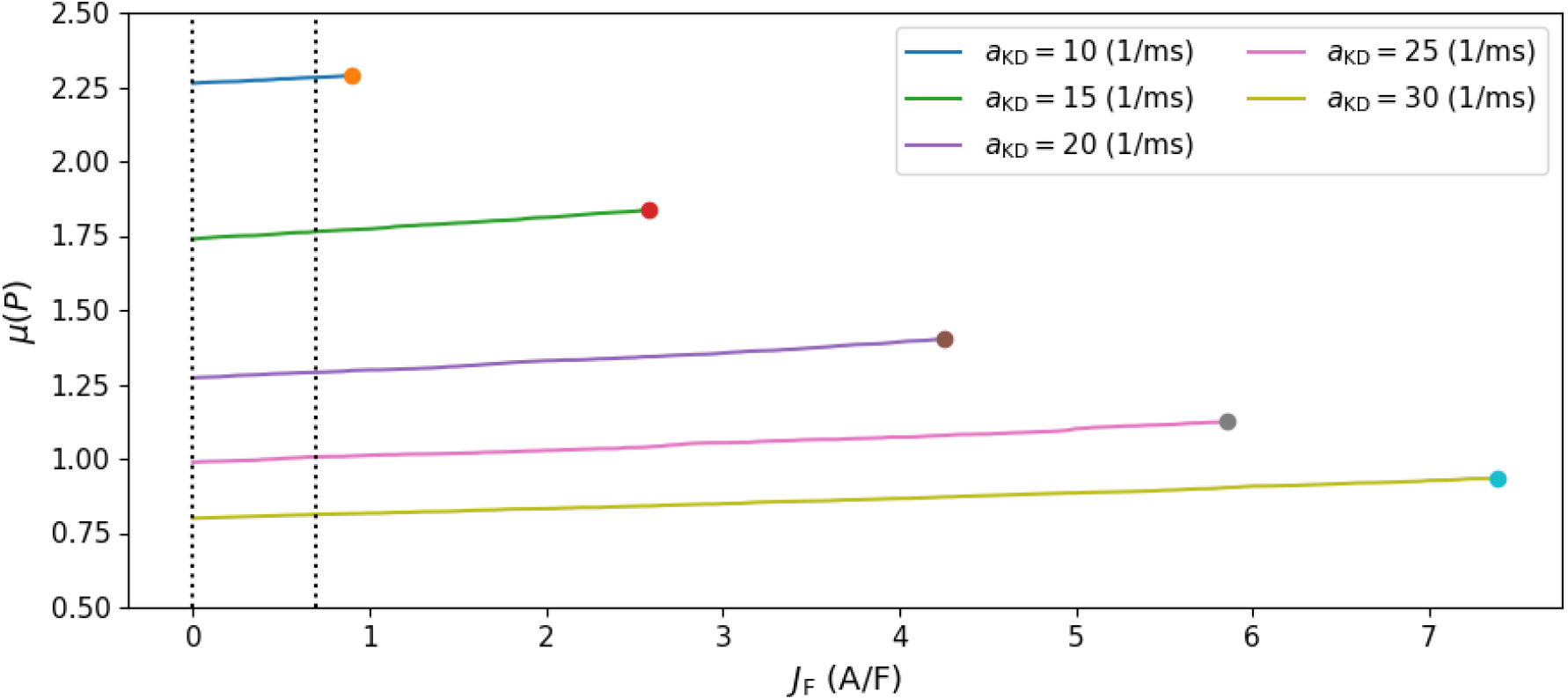
Changes in *µ*(*U*_1_) for electrophysiological phenotypes *a*_*K*_ ∈ {10, 15, 20, 25, 30 } with *J*_F_ ∈ [0, 0.75] A/F. The rheobase for each profile is marked with a dot, the upper limit for the (normalized currents), for each value of *a*_KD_.

The measure *µ* can also be used to compare the excitability of model cells representing different electrophysiological phenotypes. For instance, is it possible to decide with precision whether neurons become more or less excitable by expressing more or less potassium channels, for the same level of external stimulation?

In general terms, consider two excitable dynamical systems φ_*p*_ and φ_*q*_ with regions of excitability *U*_1*p*_ and *U*_1*q*_ in the same domain *D*, respectively associated to two parameter sets *p* and *q*. Then the system φ_*p*_ is less excitable than φ_*q*_ if *µ* (*U*_1*p*_) *< µ* (*U*_1*q*_). Using this principle, it is possible to show how the excitability decreases for electrophysiological profiles with increasing numbers of K^+^channels for *J*_*F*_ = 0 (normalized areas in Fig. 6).

#### 3.5.2 The role played by Nernst potentials

For instance, in the examples shown above *µ* (*U*_110,0_) *>* · · · *> µ* (*U*_130,0_) (Fig. 6 left-most points). So a neuron with less K^+^channels (*e*.*g. a*_KD_ = 10) is more excitable in comparison with a model neuron with more K^+^channels (*e*.*g. a*_KD_ = 30), assuming a common input current below the rheobase in both neurons, and keeping all other biophysical properties the same (Fig. 6). In general, if *a*_KD_ ∈ {*m, n*}, with *m < n*, then *µ*(*U*_1*m,J*__F_ ) *>* · · · *> µ*(*U*_1*n,J*__F_ ) for any *J*_F_ smaller than the smaller of the two rheobases.

It is also possible to compare the sizes of the excitability regions across systems. Recall that 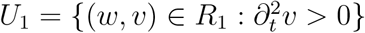. For instance, a neuron with more K^+^channels (*a*_KD_ =30) receiving a stimulus of amplitude *J*_*F*_ = 4 pA/pF is less excitable than another neuron with less K^+^channels (*a*_KD_ = 15) receiving a stimulus with amplitude *J*_*F*_ = 0.25 pA/pF (Fig. 6). However, note that monotonicity in the measured excitability with respect to *a*_KD_ is not guaranteed if *J*_*F*_ is allowed to change as well.

The Nernst potentials define the thermodynamic limits for the dynamics of *v*, establishing the physical boundaries of the phase space domain *D* = [0, 1] *×* [*v*_K_, *v*_Na_] and directly scaling the total area *A*(*D*) = *v*_Na_ − *v*_K_. To obtain a measure that could be thought of as a probability of finding an AP trajectory in the phase space, the domain could be extended by considering a larger interval for *v* that contains possible variations of *v*_K_ and *v*_Na_.

Biophysically, *v*_Na_ and *v*_K_ define the direction and magnitude of the driving forces *ψ*_KD_(*v*) ≈ *a*_KD_(*v* − *v*_K_) and *ψ*_NT_(*v*) ≈ *a*_NT_(*v* − *v*_Na_), for the voltage-gated channels, and *ψ*_NK_(*v*) ≈ *a*_NK_(*v* − *v*_NK_) for the Na^+^-K^+^ATPase^5^ (Herrera-Valdez, 2018). Varying these potentials shifts the *v*-nullcline and the boundaries of region *R*_1_. As expected, increasing the *v*_K_ decreases the driving force for K^+^, which increases the excitability of the cell (Fig. 7A). Consequently, hyperpolarizing *v*_*K*_ contracts the excitability set *U*_1_, reducing its measure *µ*(*U*_1_), which reflects on a reduced capacity for AP initiation under increased potassium driving forces. Similarly, increasing *v*_Na_ increases *µ*(*U*_1_), but now the increase in excitability is caused by an increase in the driving force for Na^+^(Fig. 7B).

**Figure 7.**
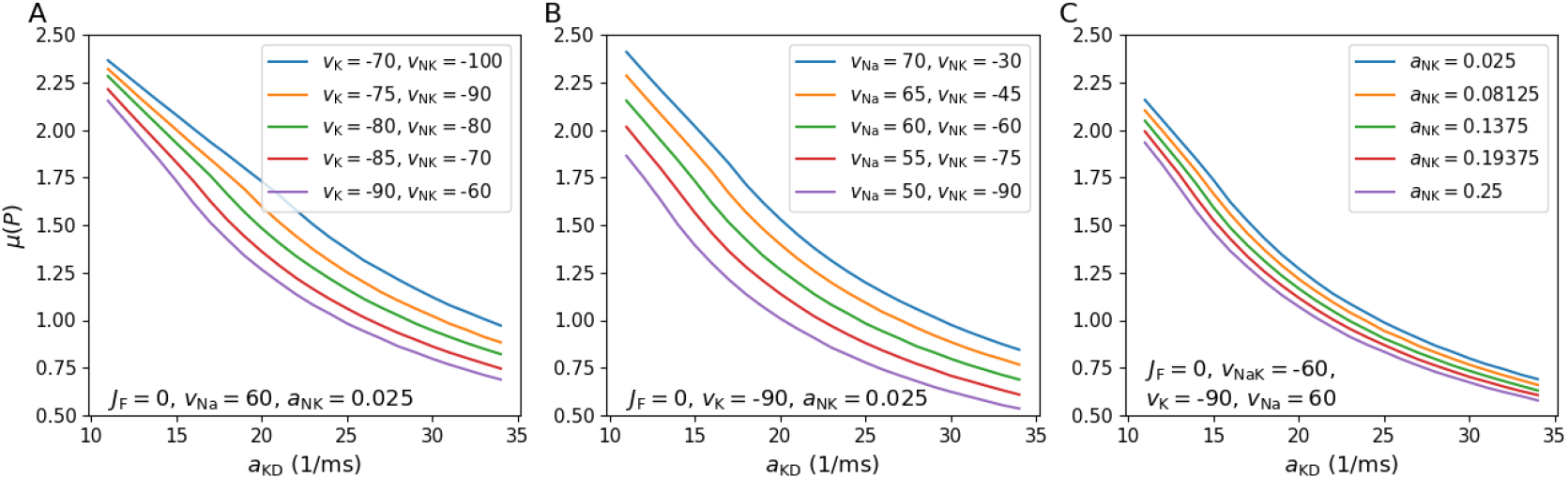
*µ*(U_1_)*fordifferentelectrophysiologicalphenotypes*,a_*K*_ ∈ {11, 35} and *J*_F_ = 0. (A)Varyng *v*_K_ ∈ {90, 85, 80, 75, 70} mV, with *a*_NK_ = 0.025 (1/ms) and *v*_Na_ =60 mV. (B)Varying *v*_Na_ ∈ {50, 55, 60, 65, 70} with *a*_NK_ =0.025 (1/ms) and *v*_K_ =-90 mV. (C) Changing *a*_NK_, for *v*_K_ =-90 mV, *v*_Na_ =60 mV, and *v*_NK_ =-60 mV, with *J*_*F*_ = 0.

Thus, decreasing the driving force for the K^+^outward current or increasing the driving force for the Na^+^inward current increase the measures of excitability in the respective dynamical systems.

#### 3.5.3 The Role of the Electrogenic Na^+^-K^+^ ATPase

The active transport of ions mediated by the Na^+^-K^+^ ATPase is modeled as a voltage-dependent pump current *J*_NK_(*v*), parameterized by a rate amplitude *a*_NK_ and its reversal potential *v*_NK_. If *v > v*_NK_, the pump generates a steady hyperpolarizing current that is essential for maintaining the resting membrane potential.

Varying the pump parameters directly impacts the system’s threshold and excitability

- The effects of changing the **pump reversal potential** *v*_NK_ on the size of *U*_1_ are non-trivial. *Depolarizing v*_NK_ by hyperpolarizing *v*_K_ (while all other parameters remain constant), increases the area of *D*, shifts the resting fixed point (*w*_∗_, *v*_∗_) to higher potentials, and shrinks the region *U*_1_. (Fig. 7A). In contrast, *depolarizing v*_NK_ by increasing *v*_Na_ (maintaining all other parameters fixed), increases the area of *D*, but decreases the range for the active outward pump current, and increases the size of *U*_1_, thus increasing the excitability of the neuron (Fig. 7B).
- Increasing the **pump rate (***a*_NK_) enhances the active hyperpolarizing outward current when *v > v*_NK_, shifting the *v*-nullcline upward in the phase plane. This active suppression counteracts weak inward depolarizing currents in the subthreshold range. This shifts the lift-off threshold *v*_LO_ to higher voltages and contracts the area of *U*_1_, thus decreasing *µ*(*U*_1_) (Fig. 7C).

### 3.6 Analytical curvature in higher dimensional models of membrane potential

Consider a more general model of neuronal membrane dynamics in which *v* ∈ (*v*_*m*_, *v*_*M*_ ) (in mV) representing the transmembrane potential, complementary variables for ion-channel gating and possibly concentrations of molecules of interest represented by a vector

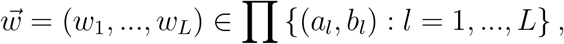

and a vector 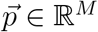 of parameters (Best et al., 2007; Rasmusson et al., 1990a; Wilson et al., 2004). Assume the dynamics are given by

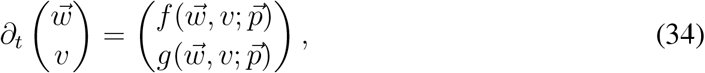

with smooth functions *f* and *g*. If we assume further that *f* and *g* are derived from biophysical principles, *f* can be thought of as a non-linear function representing the difference between a possibly time-dependent input current *J*_F_ and the sum of voltage-dependent transmembrane currents, all normalized by the membrane capacitance^6^.

*Peaks and Inflection points*. Assume that vector of parameters 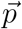 is fixed and consider the resulting dynamical system with evolution rules given by equation (34). The curvature of *v*(*t*) is described by

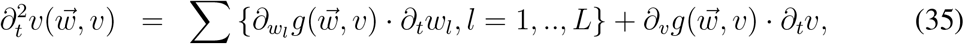

which can be calculated if *f* and *g* are smooth functions of 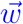 and *v*. See the Appendix, Section B for examples of explicit calculations including a case for time-dependent *J*_F_.

As found in previous sections, the 0-level curve for 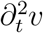 is then the set

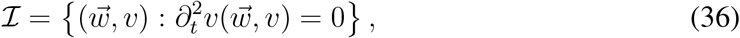

is a manifold in the 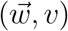-space that contains the inflection points for *v*(*t*) in the different trajectories of the system.

If 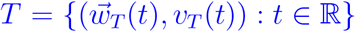 is a trajectory in phase space that contains an AP, then the AP peak is the point where *T* crosses the *v*-nullcline 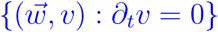, and *T ⋂ ℐ* is the set where the curvature of *v*_*T*_ (*t*) changes. The region where AP orbits cross can be characterized again in terms of the set in phase space where AP upstrokes are found, the inflection point manifold ℐ would still represent points where *v*(*t*) changes concavity, and the measures of excitability defined above can be generalized to calculate volumes in the new domain that now is a set in ℝ^*L*^ *×* ℝ.

### 3.7 Threshold manifolds in multicompartment models

Neurons are spatially distributed structures in which AP initiation sites may move depending on different factors (Bender and Trussell, 2012; Chen et al., 1997). From there, APs propagate to all the remaining compartments of the neuron, which will have different attenuating or amplifying effects on the propagating potentials. One issue of importance is how the AP thresholds change in different sites of a neuron?

To extend our single-compartment framework to a spatially explicit, a simplified multi-compartment model of *N* coupled nodes, the membrane potential *v*_*j*_ of compartment *j* can described by:

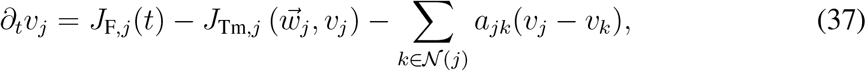

for *j* = 1, …, *n*. The transmembrane currents mediated by channels and pump proteins in compartment *j* are represented by *J*_Tm,*j*_. *J*_F,*j*_ the input current, and *a*_*jk*_ is the axial coupling conductance between compartment *j* and compartment *k*, all normalized by the compartment membrane capacitance *C*_*m*_. The set of compartments adjacent to compartment *j* is *N* (*j*).

The curvature of *v*_*j*_(*t*) changes with respect to time according to

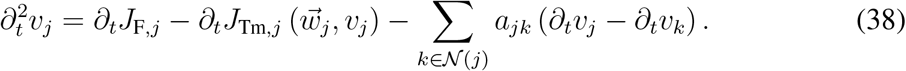

Using the chain rule, the time derivative of the local transmembrane current in compartment *j* can be calculated as in equation (35). From equation (38) it is possible to obtain the inflection points and a couple insights. In particular, at the inflection point manifold for compartment *j*,

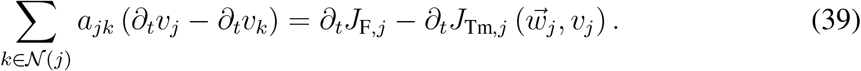

#### Active AP initiation sites

When compartment *j* initiates an action potential, its local depolarization rate ∂_*t*_*v*_*j*_ is high compared to its neighboring inactive compartments (*e*.*g*., an AP initiating in a proximal dendrite vs. soma). Consequently, ∂_*t*_*v*_*j*_ *>* ∂_*t*_*v*_*k*_. This means that axial coupling term acts as a current sink, pulling down the second derivative 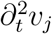 and opposing positive acceleration. This delay raises the local lift-off threshold *v*_LO,*j*_, and constitutes an electrical load that the AP initiating segment must overcome to ensure the transmission of the AP.

#### Passive downstream compartments

For a compartment *j* receiving an (excitatory) potential or a back-propagating action potential from an active neighbor *k*, the rate of depolarization in the upstream compartment is larger (∂_*t*_*v*_*k*_ *>* ∂_*t*_*v*_*j*_). The axial current acts in this case as a source of depolarizing charge, driving the local curvature positive 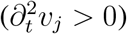 even before local voltage-gated channels have fully activated. This axial current lowers the apparent local lift-off threshold *v*_LO,*j*_, facilitating the rapid, passive acceleration of the membrane potential.

From this brief analysis, it is possible to define the AP initiation site as the location *j* where ∂_*t*_*v*_*j*_ *>* 0 and 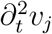 transitions to positive first, driven purely by intrinsic ionic currents (∂_*t*_*J*_Tm,*j*_ *<* 0) rather than axial source currents.

## 4 Previous work

Mathematical analysis of membrane potentials and electrical excitability has been performed in search of theoretical descriptions of thresholds (Fitz-Hugh, 1960; Noble and Stein, 1966). One outcome from that body of work is a classification for different types of threshold phenomena proposed by Fitz-Hugh (1955). However, no specific equations describing thresholds for APs have been obtained and further work has proposed approximations to address the issue. Of particular interest, Platkiewicz and Brette (2010) derived an equation that can be used to identify APs. The equation is derived from an exponential approximation for the voltage time course during the AP upstroke and the sodium current contribution to the upstroke velocity in APs (based, in turn, on approximations for voltage-thresholds, the activation voltages for Na currents, and empirical values for the ∂_*t*_*v*).

The impossibility of describing thresholds for voltage or current in models of neuronal membrane potential has also been discussed by other authors (Koch et al., 1995). From that work it has been argued that it is possible to determine a separatrix curve in 2-dimensional models of membrane potential, but only if the underlying dynamical system has 3 fixed points, with an attractor fixed point associated to the resting potential, a saddle, and a repulsive fixed point, with respectively higher *v*-values than the attractor (Izhikevich, 2007); the separatrix curve is actually the attracting portion of the *v*-nullcline above the saddle point. Another kind of boundary separating action potentials from other trajectories in phase space can be found in bistable systems having an attractor fixed point and an attractor limit cycle; in this case there is a repeller limit cycle surrounding the fixed point that can be thought of as a separatrix between AP-containing orbits converging to the limit cycle and non-action potential orbits (trajectories converging to the fixed point). However, the repeller cycle cannot be described analytically in biophysical models.

A classification for different excitability types was proposed by (Hodgkin, 1948). The classification distinguishes qualitatively different types of transitions between rest and repetitive firing as a function of the input current in the relationship between current and firing frequency (I-F). Type-I excitability is described as I-F curves with “continuous” profiles, whereas type-II excitability is related to a “discontinuous” I-F curve. It is worth remark that, from a mathematical stand point, the transitions between rest and repetitive firing in current-clamp experiments occur at bifurcations. Such transitions always involve the emergence of an attractor limit cycle for a large enough input current (the bifurcation parameter), which by definition has an intrinsic oscillation frequency that cannot be arbitrarily close to 0. That means that there is no such thing as a continuous I-F curve as described above. One additional issue is that the effect of increasing the stimulus amplitude is to change the number or the types of fixed points, and their attractivity (Herrera-Valdez, 2012, 2018). In addition, electrically excitable cells may display APs in response to stimulation that is far from the levels required for repetitive firing *in vivo* (Fellous et al., 2003; Kuhn et al., 2004; Platkiewicz and Brette, 2010; Rudolph and Destexhe, 2003). It is therefore important to consider the thresholding problem in regimes defined by different stimulus amplitudes, well below the rheobase.

## 5 Discussion

This work establishes a mathematically rigorous, geometrically grounded framework to characterize action potentials (APs) and cellular excitability without relying on localized bifurcation approximations or empirical thresholds. By focusing on the intrinsic geometry of membrane potential trajectories, it is shown that the changes in concavity during the AP upstroke provide a precise, continuous, and biophysically conserved definition of thresholds and excitability.

The motivation for this work originated from realizing that there are no formal definitions of excitability, and that defining excitability in terms of APs required first to be able to identify APs unequivocally. Since experimental work yields only time series of membrane potential, the AP identification criteria should work only for time series. The AP identification was achieved by noticing that APs display an upward concavity during the upstrokes. Once the identification was implemented and confirmed for time series, the second challenge was to use the same paradigm to characterize APs in ADS. The subsequent definition of excitability was framed around the idea that excitable systems should be able to produce pulses, but not always. From there, cellular electrical excitability can be thought of as the capability of producing both APs and non-APs (aside from trajectories at fixed points) depending on the initial conditions. The defining property for an excitable system given here is thence that the set of orbits of the system is bipartitioned, and such that one of the subsets contains only AP orbits, the other subset should contain orbits formed by multiple points.

### 5.1 A Unifying Geometric Framework for Thresholds and Excitability

The core of the approach lies in assessing the second time-derivative of the membrane potential 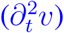, which defines the time-dependent curvature of trajectories. Specifically, the idea is to search for changes in curvature, which means looking for inflection points. In time series of membrane potential, the sampling rate of the series can hinder the accuracy of the approach. However, it is possible to define first and second order quotients of differences to calculate approximations to the first and second derivatives of *v* with respect to time. In any smooth, autonomous model, the zero-level manifold of this derivative, 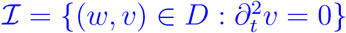, contains the inflection points where trajectories undergo concavity transitions in time.

Since AP upstrokes occur where both the membrane potential and recovery variables are increasing, the restriction of ℐto the region *R*_1_ yields the threshold manifold ℐ_*θ*_ = ℐ∩ *R*_1_, that naturally splits into two functionally distinct segments:

1. **The Lift-Off Manifold (***I*_**LO**_) containing the points of “kinetic commitment” where the upstroke of *v*(*t*) transitions from concave-down to concave-up, starting the self-amplifying acceleration of the AP.
2. **The Maximum Rate Manifold (***I*_**MR**_) formed by points where the upstroke reaches its top rate of depolarization (∂_*t*_*v*), transitioning from concave-up to concave-down as it approaches the AP peak.

Unlike prior formulations that are tied to specific bifurcation configurations or local linearizations near saddle-node points, this segmentation of ℐ_*θ*_ is defined for a wide variety of models (as long as the dynamics are smooth), analytically explicit, and robust to changes in the number of fixed points in the underlying dynamical system.

#### It is possible to identify boundaries that separate AP and non-AP orbits

In both systems φ_13,0_ (three fixed point) and φ_26,0_ (one fixed point), there is a separation of non-AP orbits and AP orbits within *R*_1_ that is *not* crossed by trajectories. If the set of non-AP trajectories starting in *R*_1_ is called *N*_1_, the separation is the boundary of *N*_1_ ^7^ (Figs. 3 and 4).

In excitable systems with three fixed points like φ_13,0_, there is another boundary between AP and non-AP orbits, namely, the segment of the *v*-nullcline connecting the saddle and the repulsive node (Figs. 3A and 4B).

### 5.2 “Excitable” vs. “Excited”

The author is not aware of any formal definitions of membrane excitability, or excitability in other contexts, only classifications (*e*.*g*. Type I-II, discussed in Section 4). A formal definition requires a clear criterion. In this case defining excitability as the capability of producing APs and other non-AP trajectories, requires, in turn, specifying what is an AP. The geometric formulation presented here allows formalizing the concept of cellular excitability in terms of a partition of phase-space trajectories into two distinct subsets: those that cross the threshold manifold *I*_*θ*_ (AP orbits) and those that do not (non-trivial non-AP orbits). The size of the region *U*_1_ is a proxy of the likelihood of finding an AP in *D*. The distinction between excitable and excited systems is a consequence of the proposed definition, but it also makes sense semantically: already excited means already producing APs, not just capable of producing them.

Notice that bistable systems in which one attractor is a fixed point, and one attractor is a limit cycle are excitable under this criteria.

This definition brings up simple but profound distinctions of physiological conditions relating to repetitive firing and oscillatory systems. First, dynamical systems possessing a unique limit cycle attractor (such as pacemaking cardiomyocytes or neurons in continuous repetitive firing) are not excitable; they are called **excited** instead. The reason is that the non-trivial trajectories in these systems converge asymptotically to the same periodic orbit, so all the multi-point trajectories contain APs. Second, when a system is driven beyond its repetitive firing regime into a **depolarization block**, the limit cycle eventually disappears and at some point all trajectories converge to a single attractor fixed point with an elevated *v*-coordinate. In this regime, the system is no longer capable of producing two distinct types of trajectories, representing a complete loss of excitability.

### 5.3 Biophysics and the excitability measure *µ*

Excitability requires the existence of a region 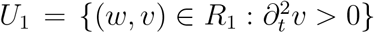 that can be found analytically and measured. Simply stated, *U*_1_ is a region of the phase space containing only upstrokes of trajectories that display a concave-up *v*(*t*). In fact, *U*_1_ contains trajectory segments belonging only to APs, and contains them all. The threshold manifold *I*_*θ*_ is the boundary of *U*_1_, and both can be found using an analytical criteria. As a consequence, the presence of a set like *U*_1_ can be used to classify the system as an excitable dynamical system, and its measure can be thought of as a multiple of the probability of finding an AP trajectory in the phase space (which could also be calculated dividing the area of *U*_1_ by the area of *D*). The measure proposed here focuses on the geometry of the phase space, and complements measures like firing frequency, delay to first spike, and others, which are concerned more with time-related properties of APs, typically obtained in contexts involving responses to a stimulus or within a context.

To quantitatively compare electrical excitability across diverse physiological states and stimulus conditions, a measure quantifying the area of the excitability region *U*_1_ is introduced. The measurements obtained from different dynamical systems can be compared because they are in the same units, and the spaces where they are defined can be assumed as common. The comparisons between the measured excitability in dynamical systems presented here agree with the biological intuition that one could use to advance conjectures about the effects of changing K-channel activation, Na-channel inactivation, the input current, and the Nernst potentials for Na^+^and K^+^, at least. Since the calculations for curvature yield expressions with all the parameters of the original models, it is not difficult to assess the progression of ℐ_*θ*_ or *U*_1_ as a function of any of the parameters in a model.

#### 5.3.1 Channel expression and kinetics

Modifying the density of voltage-gated potassium channels (*e*.*g*., varying *a*_KD_) alters the size of *U*_1_. For instance, a membrane with a lower potassium channel density (*a*_KD_ = 13) exhibits a significantly larger *µ*(*P*) than one with a higher density (*a*_KD_ = 26), showing that increasing potassium channel expression restricts excitability (Fig. 5). This occurs because K-channel activation increases the lift-off threshold, *v*_LO_, on ℐ_LO_ while decreasing the threshold for maximum depolarization rate, *v*_MR_, on ℐ_MR_, contracting the region *U*_1_ where 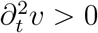.

The channel kinetics for a fixed profile of channel expression show that the voltage profile in *I*_LO_ and *I*_MR_ are respectively increasing and decreasing as functions of *w*. This means that larger proportions of K-channel activation (alternatively, more Na-channel inactivation) yield higher lift-off threshold potentials, and lower maximum-rate threshold potentials, respectively.

### 5.4 Biological Generality Beyond Neurons

A major strength of this concavity-based approach is its mathematical and physiological universality. The geometric criteria presented here transcends neurobiology, and the same analytical tools can be applied to describe action potential thresholds in other types of excitable cells. Of possible interest, further work could be done to study properties of (a) *cardiomyocytes and endocrine cells* where specialized pumps and channels regulate cardiac and pacemaking activity, or *α*- and *β*-cells that secrete glucagon and insulin, respectively; (b) *astroglia and striated muscle fibers* in which slow active propagation or calcium release depends on threshold-like membrane responses; (c) *plant cells*, in particular vascular plants, that possess excitable membranes that use APs to communicate across long distances (*e*.*g*., AP in *Mimosa pudica*); and giant algal cells (*e*.*g*., *Chara* or *Nitella*), which exhibit classic “all-or-none” electrical pulses driven by transient chloride and potassium efflux.

The method presented here remains invariant to the specific biophysical representation of the transmembrane fluxes and channel or pump kinetics, making it equally applicable across the thermodynamical model (Herrera-Valdez, 2018), conductance-based models (Av-Ron et al., 1991; Fohlmeister and Miller, 1997; Hodgkin and Huxley, 1952; Rinzel, 1985). In addition, the method is also applicable to phenomenological models like the Fitz-Hugh (1961) model (see examples in the Appendix, Section B).

### 5.5 Limitations and Future Directions

While the deterministic, autonomous framework presented here is mathematically complete, applying it to real-world physiological conditions introduces exciting avenues for expansion:

#### 5.5.1 Stochastic Membrane Fluctuations

Real cells operate in noise-driven regimes where synaptic background activity or channel noise induces random membrane potential fluctuations. Under such non-autonomous conditions, the size and shape of the excitability region *U*_1_ become time-varying. Therefore, it becomes is necessary to consider a time-varying *U*_1_(*t*) over sample times *t*_0_, …, *t*_*N*_ . By analyzing time-series representing the excitability region *U*_1*i*_ = *U*_1_(*t*_*i*_), at time *t*_*i*_ it would be possible to obtain a sequence of excitability measures *µ*_*i*_ = *µ*(*P*_*i*_), *i* = 0, …, *N*, to quantify how fluctuating synaptic drives or current stimuli dynamically modify the probability of AP initiation over time.

#### 5.5.2 Spatial compartmentalization and multicompartment formulation

A further limitation of the present study is its focus on single-compartment dynamics rather than spatially explicit models of membrane potential, which are necessary to capture the complex dendritic and axonal geometries of biological neurons. Experiments and models show that sites of initiation of action potentials in neurons may move depending on factors that include the synaptic load on the dendrites (Bender and Trussell, 2012; Chen et al., 1997). One possible extension of interest would be to calculate second time-derivatives of *v* in multi-compartment models to detect and analyze AP initiation sites (equation (38)). In essence, the calculations would not change, but now there would be one threshold manifold per compartment. Different regions in a spatially explicit model should have different threshold profiles. Since the biophysical attributes captured in the equations are preserved in the calculations for curvature, it would be reasonable to expect different compartments to display qualitatively different thresholds in terms of their biophysical and geometrical properties. Current work in this direction is underway.

The formulation in equation (38) reveals the influence of axial coupling on local upstroke curvature (discussed in Section 3.7). By calculating the spatial profiles of curvature with respect to time, researchers can mathematically map and identify the exact compartment where the active “kinetic commitment” occurs.

### 5.6 Conclusion

The concavity criteria proposed here moves beyond static bifurcation analysis to a dynamic interpretation of the phase space, and provides a general, mathematically rigorous criterion to assess excitability in the dynamical systems that compose a model. Therefore, the analytical description of excitability presented here unifies the theory of excitable dynamical systems for continuous models by providing a tool to compare excitability across different dynamical systems. Further, the work presented here provides connections with experimental data, bridging the theoretical multivariate descriptions of systems with voltage recordings.

## Acknowledgments

This study was funded by the DGAPA-UNAM grants PAPIIT-IN228820 and PAPIME-114919. The author would like to thank the members of the Dynamics, Biophysics, and Systems Physiology Laboratory (Stephanie Escobar-Sanchez, Germán Pérez-Chavez, Carlos Andrés Gil-Gómez, and Olyn Castañeda Ramírez for discussing the first ideas behind this article, and to Hector Chaparro-Reza, Vania Victoria Villegas-Martinez, Roberto García-Medina, Carlos Gil Gómez, and Lionardo Truqui for checking some of the calculations).

This work is dedicated to Eréndira Valdez Coiro and Ignacio Arieli Herrera Heredia.

## Appendix

### A Methods

#### Nullclines and fixed points

From equations (15)-(18), the points (*w, v*) in the *w*-nullcline is satisfy either *w* = 0, or

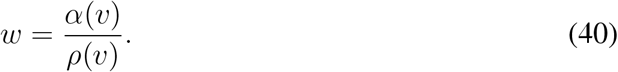

whereas the points (*w, v*) in the *v*-nullcline satisfy

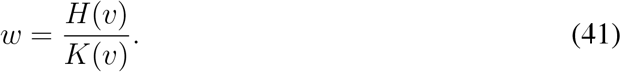

The *fixed* points (*w*_∗_, *v*_∗_) of the system *F* (Fig. 3A1,B1), are the points in the intersection of the two nullclines. One way to obtain the values of *J*_*F*_ that with the corresponding values of *v*_∗_ is to use

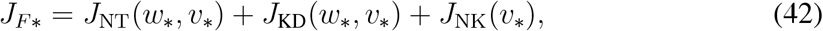

with *w*_∗_ = {1 + exp [*s*_*w*_ (*v*_*w*_ − *v*)]}^−1^.

The type and attractivity of each fixed point (*w*_∗_, *v*_∗_) can then be determined analytically by calculating the Jacobian matrix

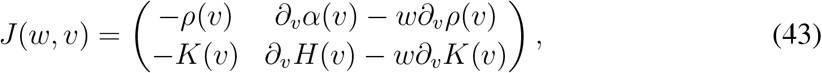

and evaluating at (*w*_∗_, *v*_∗_) to obtain the trace, determinant and discriminant of the characteristic polynomial (Strogatz, 1994).

#### Curvature and inflection points

The inflection points can be calculated using equations (**??**), (16), and (15). The idea is to take into account that *v* and *w* are functions of *t* and use the chain rule for differentiation. The set ℐ of inflection points (*w, v*) for the trajectories of the system satisfies

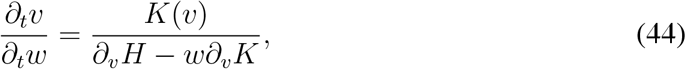

which can be written explicitly in terms of the original expressions that give *H*(*v*), *K*(*v*), *α*(*v*), and *ρ*(*v*) (equations (16)-(15)).

### B Curvature calculations in 2D models

#### B.1 Av-Ron et al. (1991) minimal biophysical model (similar to the Hodgkin and Huxley (1952) model but 2-D)

The model proposed by Av-Ron et al. is very similar to the Hodgkin and Huxley model, but takes into account that the voltage-dependent activation for Na is very fast, and the voltage-dependent inactivation for Na varies as a linear function of the voltage-dependent activation for the K-current. For details on the reduction see (Av-Ron et al., 1991; Herrera-Valdez and Lega, 2011; Rinzel, 1985). Explicitly,

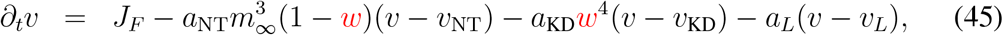

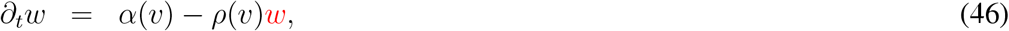

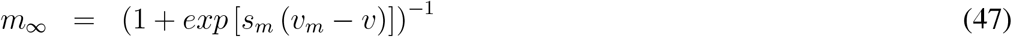

Let

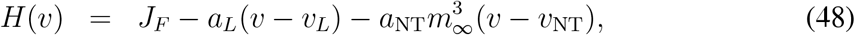

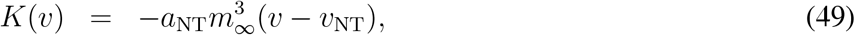

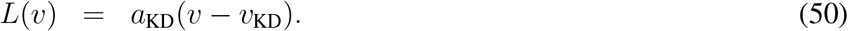

Then

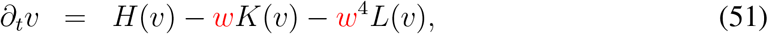

which means

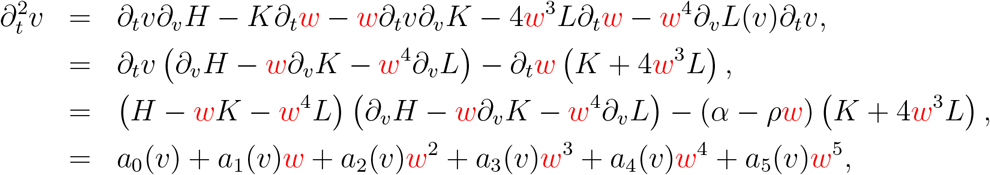

with

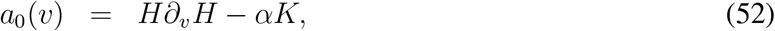

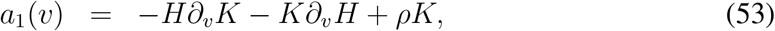

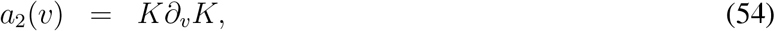

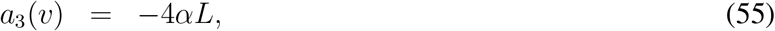

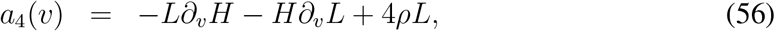

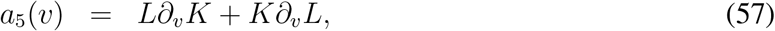

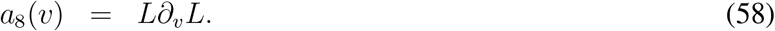

As before, the curve ℐ can be found as in expression (26).

#### B.2 Fitz-Hugh model

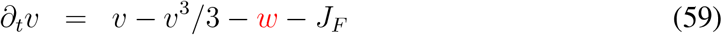

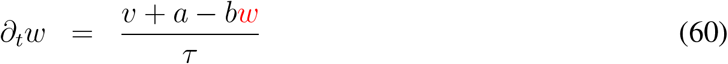

Example: *a* = 0.7, *b* = 0.8, *τ* = 13, *J*_*F*_ = 0.01.

Nullclines given by functions

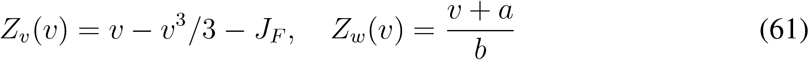

Curvature:

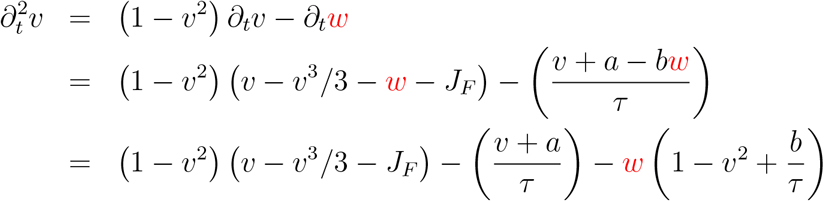

Inflection points:

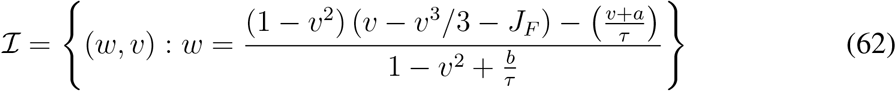

#### B.3 Up-Down model

This is a minimal model for the membrane potential involving two large currents inspired by the work of Herrera-Valdez and Lega (2011). The “Up”-current increases *v* (*e*.*g*. Na-current), the “Down” current provides negative feedback to *v* (*e*.*g*. K-current). The equations can be written explicitly in conductance-based form as:

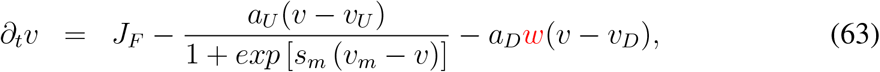

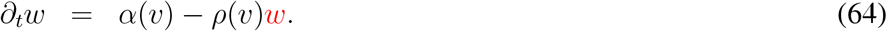

Let

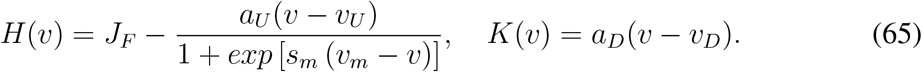

Nullclines given by functions

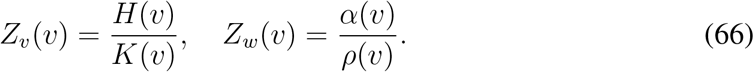

The curve ℐ can be found as in expression (26), which has a closed form solution that can be obtained using the quadratic formula.

In all cases, the threshold manifold is the set of inflection points within the region *R*_1_, such that

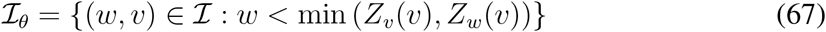

#### B.4 Time-dependent input current

If the input current is a smooth time-dependent function, it is still possible to use equation (**??**) to calculate the curvature and from there, the inflection points. In this case, the *v*-nullcline would move with time, and the inflection point manifold would also change with respect to time (the system is no longer autonomous). To illustrate how the calculation could be done, consider the Up-Down model and a time-dependent input current *J*_*F*_ (*t*) (normalized by membrane capacitance). The equation for *v* would be

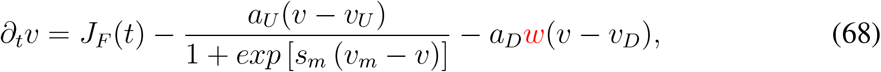

which can be rewritten as

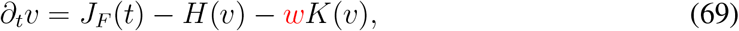

with

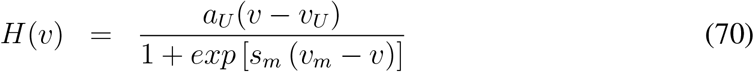

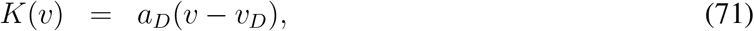

The *v*-nullcline would be given by the function

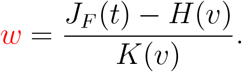

The curvature would change according to,

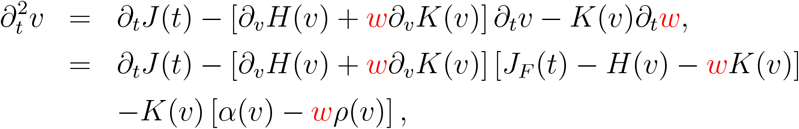

which yields inflection points when

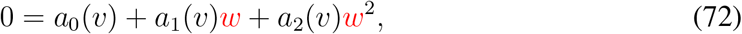

In that case the coefficients transform into

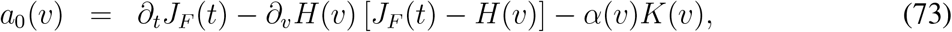

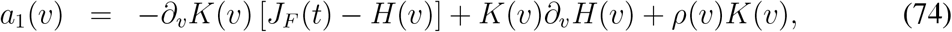

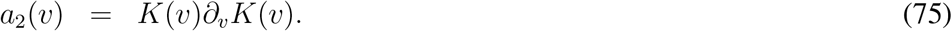

For instance, if the current stimulus was a ramp with slope *ε >* 0, then the function defining the *v*-nullcline would change to

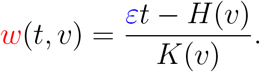

and the coefficients *a*_0_ and *a*_1_ would change to

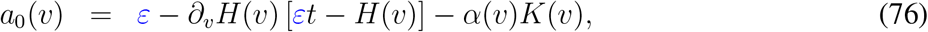

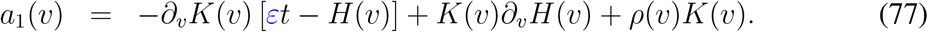

### C Autonomous 2-D dynamical systems modelling the membrane potential of cells have at least one attractor

Consider 2-D, continuous, models of the membrane potential with smooth evolution rules. For biophysical models, consider the Hodgkin and Huxley (1952) model based on equivalent circuits and conductance-based currents, or more general models not derived from equivalent circuits (Herrera-Valdez, 2020) that model pump and channel-mediated currents as well (Herrera-Valdez, 2018). For phenomenological models, consider the Fitz-Hugh (1961) model. Do these autonomous, 2-D continuous dynamical systems have at least one attractor?

The short answer is yes. Any dynamical system φ as the ones mentioned above has at least one attractor, that is either a fixed point or a limit cycle. The bounded, 2-D non-linear dynamics in such systems system, work together to ensure an attractor exists.

For more details, note first that the domain in such models will be a subset of a metric space (ℝ ^2^), and it will also be closed and bounded if the model assumes realistic biophysical conditions. Therefore, the domain is a compact subset of ℝ^2^.

#### Forward Invariance

The guarantee relies on *D* being forward-invariant (a trapping region). *D* is naturally forward-invariant because physical constraints prevent variables from growing indefinitely: in physiological conditions the voltage *v* is bounded by the Nernst reversal potentials of sodium and potassium (*v*_*K*_ ≤ *V* ≤ *v*_Na_), and the recovery variable *w* is bounded between 0 and 1. The vector field along the boundary ∂*D* points inward. If *D* was merely an arbitrarily drawn bounded box that is not forward-invariant, trajectories could flow straight out of *D* in finite time, leaving no attractor inside *D*.

#### The system has a global attractor

Under set-theoretic definitions, if *D* is compact and forward-invariant, the sequence of images under time evolution φ_*t*_(*D*) forms a nested family of compact sets. The intersection:

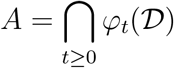

is a non-empty, compact, invariant set *A* ⊆ *K* that attracts all initial conditions in *D*. This set *A* is the global attractor of the system. Under Milnor’s measure-theoretic definition ^8^, *A* contains at least one Milnor attractor with a positive-measure basin of attraction.

#### The Poincaré-Bendixson Theorem guarantees the existence of the attractor

In a 2-D continuous autonomous system defined on a plane (ℝ^2^), dynamics are tightly constrained by topology. Note that the compact domain *D* is forward-invariant, meaning that its trajectories that start inside *D* can never leave *D*. The Poincaré-Bendixson Theorem(Guckenheimer and Holmes, 1990; Strogatz, 1994) dictates that the *ω*-limit set (*i*.*e*. the long-term destination) of every trajectory in *D* must be one of three things: (1) a fixed point where ∂_*t*_*v* = 0 = ∂_*t*_*w*); (2) a periodic orbit (limit cycle); (3) a connected union of fixed points and connecting orbits (*e*.*g*. homoclinic/heteroclinic loop).

*D* is compact (closed and bounded) and traps trajectories, the flow cannot diverge to infinity or exhibit 3D-style chaos (by the Poincaré-Bendixson theorem). Trajectories starting inside *D* must settle into at least one of these bounded structures.

#### Biological dissipativity prevents conservative dynamics

For continuous systems in general, a compact space could theoretically contain neutrally stable orbits (like nested conservative circles in a frictionless pendulum) without a proper attractor. However, cell membrane dynamics are inherently dissipative. Among other reasons, membrane models balance driving currents (ion pumps, applied current) against dissipative losses (currents mediated by channels). Thermodynamically, the phase-space volume contracts on average (∇ · *f <* 0 across large regions of *D*).

Dissipativity breaks neutral stability, so at least one of the invariant sets, say *S* inside *D* must act as a sink. Then either *S* is a point attractor, with its *v*-coordinate representing the resting membrane potential, or *S* is a limit cycle (closed orbit) attractor that represents continuous, repetitive tonic firing of APs.

### D A note on bifurcation structures and rest-repetitive firing transitions

Assuming that the system does not already exhibit repetitive spiking in the absence of current stimulation, the number of fixed points for the system depends on the ratio *a*_KD_*/a*_NT_ that can be roughly thought of as *N*_KD_*/N*_NT_, the number of K channels divided by the number of Na channels. The same happens for small enough values of *J*_*F*_ . Roughly speaking, lower *N*_KD_*/N*_NT_ ratios correspond to three fixed points and higher *N*_KD_*/N*_NT_ ratios eventually yield one fixed point. As a consequence, the system for *J*_*F*_ = 0 may have three, two, and one fixed point respectively (Fig. 3). The configuration with two fixed points is unlikely to be observed experimentally, and when studied in models it is most of the time only accessible through numerical approximations.

Another way to observe the transition between three and one fixed point is to increase the stimulus amplitude *J*_*F*_, which will eventually lead to a saddle-node bifurcation in which two fixed points will be lost as they collide before disappearing.

The transition between three and one fixed point can also be observed by changing parameters like *s*_*w*_ and *v*_*w*_, which affects the slopes of the *w*-nullcline, or the middle branch of *v*-nullcline.

For small enough *J*_*F*_ (values below the rheobase at which the rest to repetitive firing occurs), the fixed point with the smallest *v*-value is an attractor, and its *v* value corresponds to the resting potential (Herrera-Valdez, 2012; Herrera-Valdez et al., 2013).

It should be noted, however, that in both configurations, the sequence of fixed point bifurcations with respect to the parameter *J*_*F*_ includes a subsequence of bifurcations that include switching attractor nodes to attractor foci, repeller foci, attractor foci, and attractor nodes, in that order (Fig. 8). Limit cycle attractors emerge through fixed point bifurcations (e.g. saddle-node on invariant circle, or supercritical Andronov-Hopf) or through limit-cycle bifurcations (e.g. saddle-node off invariant circle, or fold) that result in bistability with one attractor focus point and an attractor limit cycle (Herrera-Valdez, 2012; Herrera-Valdez et al., 2013).

**Figure 8.**
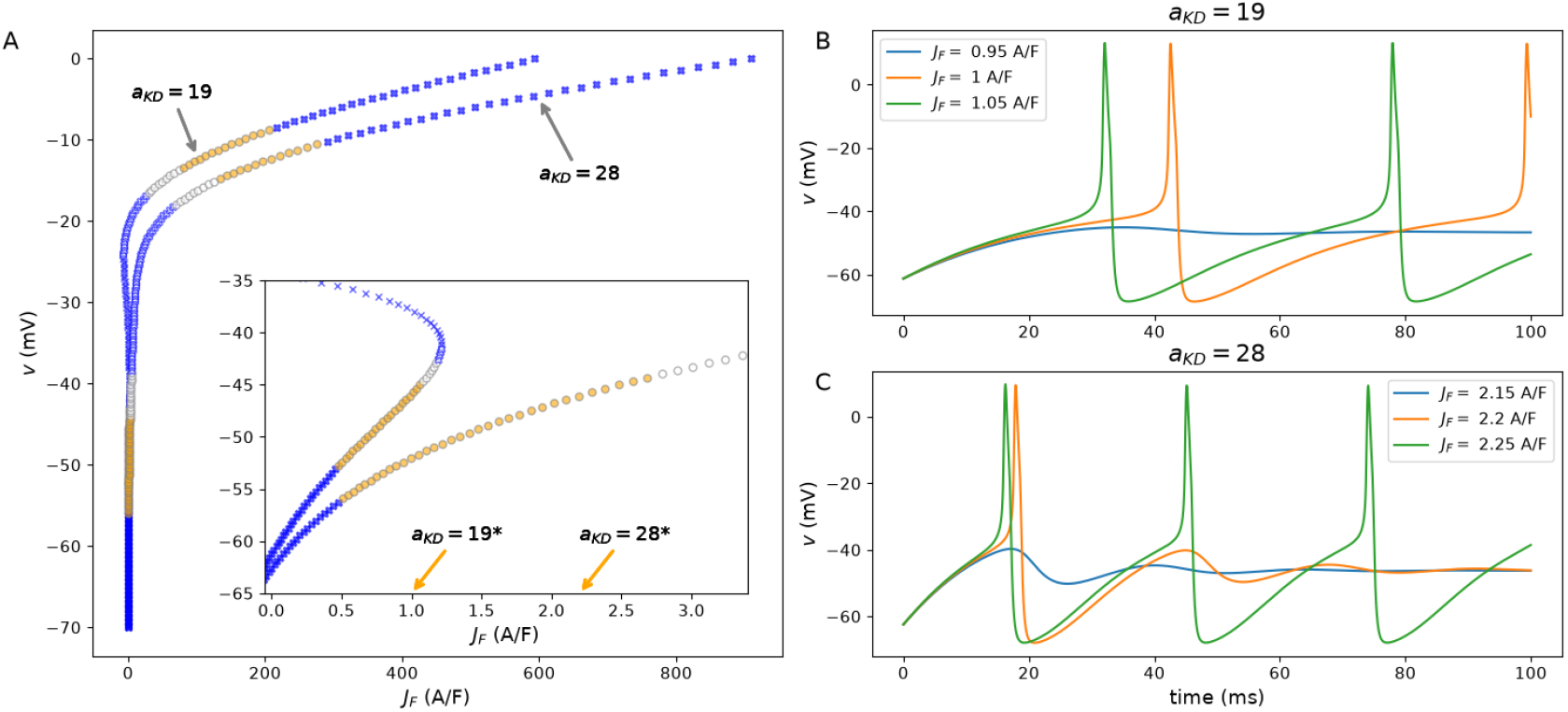
Bifurcation diagram for the two configurations of the model with *a*_KD_ = 19 1/ms and *a*_KD_ = 28 1/ms, respectively. The filled points represent attractor fixed points. Blue crosses represent attractor nodes and filled yellow circles represent attractor foci. Repulsive fixed points are represented by white-filled circles (nodes in blue crosses filled, foci in gray circles). Saddle points are represented with thin blue crosses. Orange arrows indicate the *J*_*F*_ values where the transition between rest and repetitive spiking would occur in a current-clamp experiment. B-C. Rest to repetitive spiking transitions for I-Clamps with 1 pA of resolution for the input current (*C*_*m*_ = 20 pF). The orange traces show the trace with the first AP at a resolution of 1 pA.

## Footnotes

1 The trajectories (or orbits) of any continuous dynamical system either consist of one point (start and remain at a fixed point), or start elsewhere and pass through multiple points. Trajectories of the latter kind are referred to as non-trivial.

2 Assume that *D* is a metric space. A continuous dynamical system is as function *φ* : ℝ*× D* → *D* such that (i) *φ*(0, *x*_0_) = *x*_0_ for all *x*_0_ ∈ *D* (initial condition property), and (ii) *φ*(*s* + *t, x*_0_) = *φ*(*s, φ*(*t, x*_0_) (group property). If an ordinary, autonomous differential equation defined on has solutions for all initial conditions in ∈ *D* satisfying (i) and (ii), then the mapping describing all such solutions it can be thought of as an autonomous dynamical system (Kloeden and Rasmussen, 2011).

3 The thermodynamical model for transmembrane transport is a unifying biophysical framework derived from non-equilibrium thermodynamics that allows modeling transmembrane fluxes mediated by pumps and channel with a common mathematical formulation.

4 Formally defined here as inf *J*_*F*_ for which the model displays repetitive spiking starting at the point (*w*_∗_, *v*_∗_) that corresponds to *J*_*F*_ = 0, as in current clamp experiments.

5 The pump is electrogenic, typically exchanging 3 intracellular Na^+^ ions for 2 extracellular K^+^ ions after ATP hydrolisation, so *v*_NK_ = *v*_ATP_ + 3*v*_Na_ − 2*v*_K_.

6 More precisely, the voltage-dependent change in the charge at the water membrane interface (in pF), usually regarded as the membrane capacitance, (Cole and Hodgkin, 1939; Herrera-Valdez, 2020)

7 The boundary of a set *B* is the set of points in the closure of *B* that is not in the interior of *B*. For instance, for (*a, b*], the interior is (*a, b*), the boundary is the set {*a, b*}.

8 John Milnor proposed a measure-theoretic definition in 1985 to fix the limitations of classical topological definitions: A closed invariant set *A* is a *Milnor attractor* if its basin of attraction: *B*(*A*) = {*x* ∈ *K* : *ω*(*x*) ⊆ *A*} has positive measure, and no strictly smaller subset of *A* attracts almost all points in *B*(*A*).

